# Development of the cognitive energy landscape from infancy to adolescence

**DOI:** 10.64898/2026.06.17.732977

**Authors:** Huili Sun, Isabella Stallworthy, Dustin Scheinost, Dani S. Bassett

## Abstract

Brain networks mature in a hierarchical sequence that parallels the emergence of cognitive functions. However, precisely how brain structural maturation supports the ordered development of cognitive functions remains largely unknown. Here, leveraging network control theory and 3712 developmental diffusion magnetic resonance imaging scans, we investigate how the brain’s structural effort to activate cognitive states — quantified as control energy — changes from infancy through adolescence. A total of 100 cognitive states were defined as meta-analytic activation maps from NeuroSynth, prioritized by their frequency in major neurodevelopmental behavioral assessments. We show that the control energy to drive most cognitive tasks decreases during development (for 96 out of 100 cognitive states). Ages to achieve optimal energy efficiency for each state concentrate around school age and late adolescence, whereas social and perceptual functions reach efficiency earlier (mean optimal age = 100.2 months) than higher-order cognitive functions (mean optimal age = 205.5 months). Further, we estimated the influence of molecular-level neurodevelopmental events on control energy by coupling control inputs to each event’s gene expression profile. We find that such influences vary in both temporal breadth and cognitive scope. Prenatal events (neuron differentiation and migration) exert effects mostly in infancy, while the prolonged process of myelination shapes the energy landscape across all developmental periods and the widest range of cognitive domains. Moreover, the transition energy architecture remains stable across development but becomes progressively modularized, such that transitions within the same category of cognitive states become increasingly favored. Together, these findings provide a comprehensive growth chart of how brain structural maturation supports the hierarchical emergence of cognitive abilities across early life, and establish a normative framework that enables systematic approaches to activate targeted brain circuits and facilitate selective cognitive transitions.

## Introduction

Brain networks develop in a hierarchical sequence, maturing from sensory and motor areas to higher-order cognitive association regions, to support various cognitive functions^1–3^. This maturation process is made possible by the prolonged development of white matter^4^. The complex pattern of white matter tracts undergoes dramatic changes from infancy to adulthood, progressively increasing the efficiency of neural communication^5,6^. The maturation of long-range tracts then facilitates the integration of specialized regions into large-scale networks, such as the default mode, frontoparietal, and cingulo-opercular networks^7,8^. The order in which brain networks mature broadly parallels the order in which cognitive functions develop: basic perceptual and motor skills develop first, followed by the later-emerging abilities for language, decision-making, and cognitive controls^9,10^. Although prior studies have examined how the development of brain networks supports their corresponding behaviors, a comprehensive and quantitative map linking the maturation of brain structural networks to the unfolding of specific cognitive functions remains elusive.

One promising approach to closing this gap is Network Control Theory (NCT), which provides a quantitative framework for linking brain structural connectivity to cognitive dynamics^11,12^. Rooted in control engineering^13^, NCT treats the brain as a networked dynamical system in which white matter connectivity determines how neural activity propagates and evolves over time^14^. To stimulate state changes in any system, certain input is needed to select parts, and this input energy can be quantified as control energy ^15^. For the dynamic brain system, the control energy tracks the cognitive effort required to switch between tasks^16,17^ and can be used to study external stimulation from the environment (i.e., visual and auditory stimuli)^18^ and assess clinical treatments (i.e., TMS and deep brain stimulation)^19,20^.

NCT has revealed meaningful relationships between structural organization and the energetic cost of activating cognitive states during development^21,22^. Prior work demonstrated that the energy cost to activate the frontalparietal network declines with age^23^, and that the asymmetry in transition energy along the sensory-to-association cortical axis diminishes from school age to young adulthood^24^. However, the application of NCT to understand brain development has remained limited by several methodological factors. First, existing studies relied on relatively small samples, typically around a few hundred subjects, constraining statistical power and generalizability across datasets. Second, such studies focused on narrow age windows, leaving the full developmental arc from infancy to adolescence largely uncharacterized. Third, and critically, the cognitive state spaces used in these analyses have been limited. NCT requires explicit functional activation maps as target states, and prior work has relied either on canonical functional network parcellations (e.g., the Yeo 7-network atlas)^21,23^ or on task-fMRI activation maps collected within the same neuroimaging dataset^25,26^. Both paths constrain the breadth of cognitive domains that can be examined and make it difficult to systematically evaluate control energy across the full landscape of human cognition.

Here, we address these limitations. Leveraging ∼4,000 diffusion MRI scans spanning infancy to adolescence across 11 sites, we construct individual brain structural connectomes. We defined 100 cognitive states as meta-analytic functional brain activation maps from NeuroSynth^27^, prioritized by their frequency of occurrence across major neurodevelopmental behavioral assessments. This approach enables, for the first time, a quantitative characterization of the full developmental trajectory of control energy across a broad cognitive landscape, and of the complete cognitive state-switching map, with the 100×100 matrix of pairwise transition energies between cognitive states. We further investigate the molecular underpinnings of this landscape by weighting regional control inputs by gene expression profiles for six neurodevelopmental processes derived from the Allen Human Brain Atlas^28^. Together, this approach offers a unified, energy-based account of how cognitive development unfolds in order, grounding the hierarchical emergence of cognitive abilities in the structural maturation of the brain.

## Results

Leveraging network control theory, we aimed to examine the changes in control energy required to support cognitive tasks during development. With around 4000 DTI scans collected from children across 11 imaging sites, we built individual brain structural connectomes, from infancy to adolescence. Next, to identify key concepts in developmental cognitive science, we aggregated the behavioral assessments from large-scale neurodevelopmental datasets and collated 100 cognitive keywords with the highest occurrences. Then, the brain activation patterns related to these cognitive tasks were defined with NeuroSynth (Fig. 1). We bring the brain structural connectome and activation patterns together to quantify the control energy required to activate each cognitive task and transition between any two cognitive tasks, and then we investigate how these energy requirements change across the early lifespan.

**Fig. 1.**
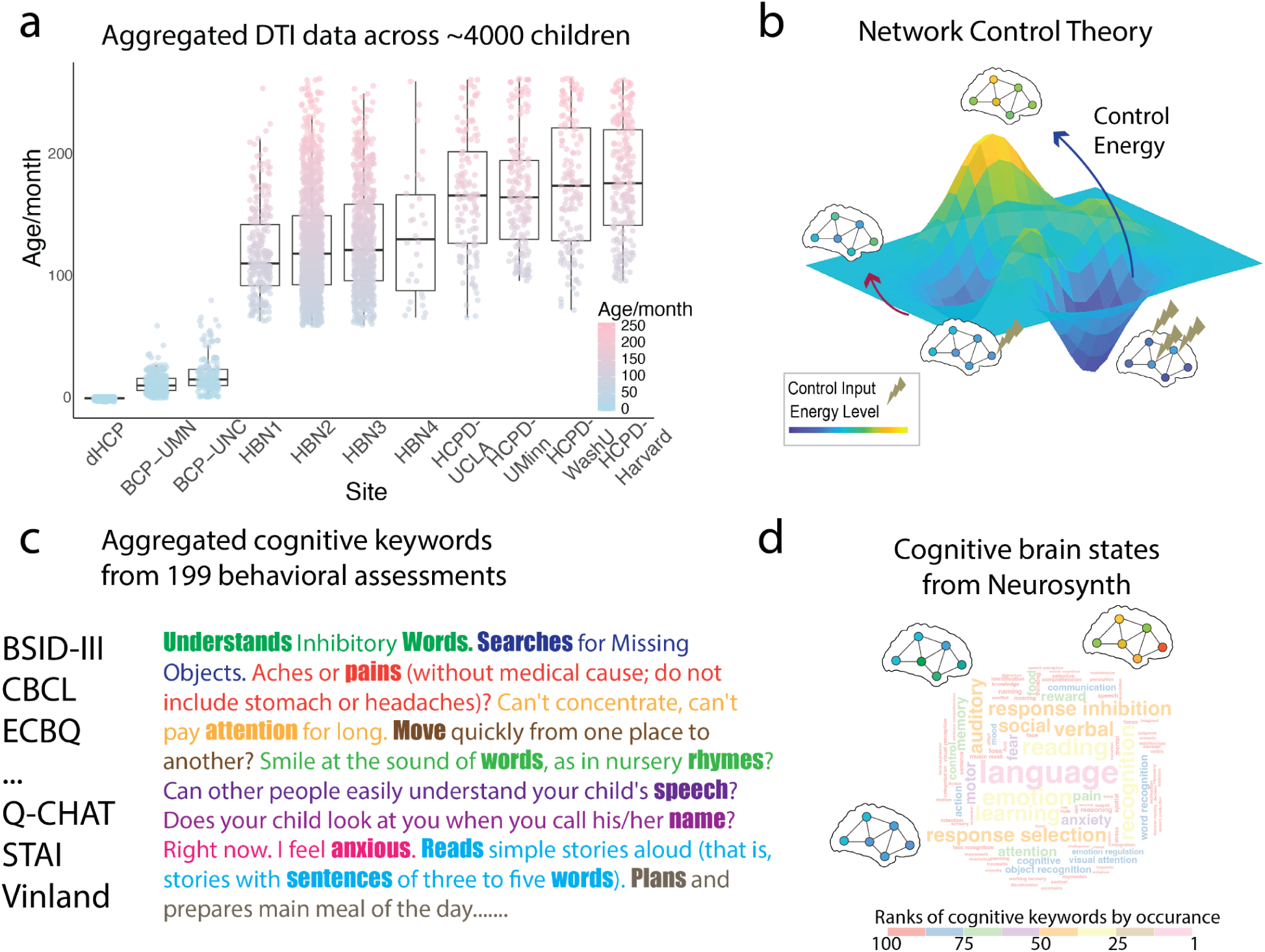
Overview of study design. **a,** Quality-controlled DTI data from around 4000 children were aggregated from four developmental neuroimaging datasets. **b,** Network control theory was used to quantify the theoretical energy cost for the brain to drive state transitions between cognitive tasks. **c,** Around 200 behavioral assessments were aggregated together. Based on the occurrences, 100 cognitive keywords were defined to be keywords of interest. **d,** Target cognitive brain states were defined using meta-analytic activation maps from the NeuroSynth database.

We first evaluated developmental changes in the control energy required to activate cognitive task-related brain states. To define these task-specific activation patterns, we used a data-driven approach to identify representative cognitive topographies. We began with the Cognitive Atlas^29^, which provides a comprehensive ontology of cognitive and behavioral terms studied in cognitive science. To better capture topographies relevant to neurodevelopment, we reweighted this vocabulary by quantifying term frequency within 119 behavioral assessments commonly used in large-scale developmental cohorts, such as the Bayley Scales of Infant Development^30^, the Child Behavior Checklist^31^, and the Vineland Adaptive Behavior Scales^32^ (Fig. S1). From this procedure, we selected the top 100 most frequently occurring cognitive keywords, spanning multiple domains, with the largest representation in language (n = 16), followed by emotion and learning (n = 13 each; Table S1). These keywords were then used as inputs to NeuroSynth to derive corresponding meta-analytic brain activation maps, which served as proxies for the brain activation patterns associated with each cognitive state in downstreaming control energy analyses.

We next quantified the control energy required for each of the 100 cognitive states across 4,000 individual structural connectomes spanning infancy to adolescence. Developmental trajectories were modeled using generalized additive models for location, scale, and shape (GAMLSS), with sex and diffusion MRI scan quality included as fixed-effect covariates and scanner site modeled as a random effect. The developmental trajectories and developmental rate curves of six cognitive tasks with top occurrences were show-cased in Fig.2a, with the rest of 100 states’ whole picture in Fig S2. All 100 GAMLSS models converged successfully, with the model fitting performance reported in Table S2.

**Fig. 2.**
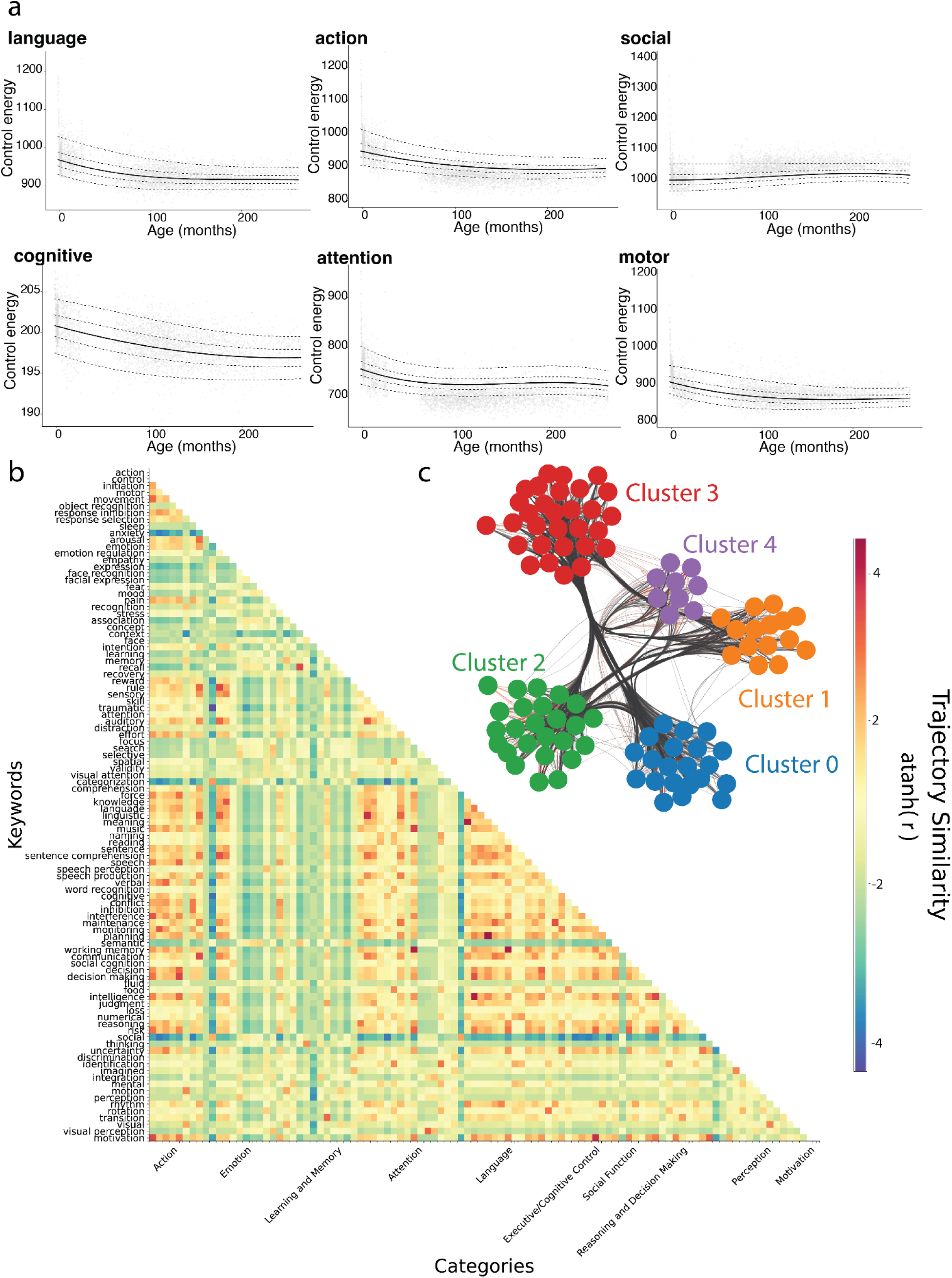
Development of control energy required to activate brain cognitive states. **a,** Developmental trajectories of control energy for six representative cognitive states with the highest occurrence. Each panel shows individual data points (gray dots) and the fitted GAMLSS growth curve (solid line) with 95% confidence intervals (dashed lines). **b,** Pairwise similarity matrix of developmental trajectories across all 100 cognitive tasks. **c,** Network visualization of the five trajectory clusters, with nodes representing individual cognitive states (color-coded by cluster membership) and edges indicating high trajectory similarity.

In the majority of trajectories (96/100), control energy decreased from infancy to adolescence, consistent with a shift toward more energy-efficient brain dynamics over development. Among the decreasing states, 36 declined monotonically throughout development, while 55 exhibited a transient recovery period — first decreasing, then briefly increasing, before declining again. This pattern was found in cognitive domains such as attention, memory, perception, and face recognition. Five states, including emotion regulation, focus, learning, motor, and social cognition, followed a U-shaped trajectory. Three states (i.e., anxiety, categorization, and sleep) showed the inverse pattern, initially increasing before declining. Finally, a single state, context, followed a non-monotonic increasing trajectory, rising, then falling, then rising again across development.

Despite the overall decrease, the shapes of the developmental trajectories are different across the cognitive landscape, raising the question of which cognitive functions are developing together. To quantify the similarity among the trajectories, we computed pairwise correlations between the mean centile curves of all 100 control energy trajectories, yielding a 100×100 similarity matrix that captures the extent to which cognitive states share common developmental patterns (Fig. 2b). We then applied the k-means clustering to these trajectories and identified five data-driven clusters (Fig. 2c; Table S3) using the elbow method (Fig. S3). Alternative clustering algorithms, including hierarchical clustering, spectral clustering, and Gaussian mixture modeling, yielded similar results (Fig. S4).

Notably, the data-driven clusters did not fully align with canonical cognitive categories. This divergence was particularly evident within the emotion category: keywords such as pain, fear, and anxiety (cluster 2) followed developmental trajectories distinct from those of emotion regulation, arousal, and stress (cluster 1). In contrast, greater alignment between clusters and canonical domains was observed elsewhere: cluster 4 showed substantial overlap with learning and memory (44% of cluster members), and cluster 5 aligned most closely with reasoning and decision-making (28% of cluster members), suggesting that these functional categories are developmentally more coherent (Fig. S5).

Developmental rate curves were calculated as the first derivative of the median centile line (Fig. 3a). With the developmental curves, we then asked when the control energy approaches its minimum (i.e., most energy-efficient) state during development. We defined this optimal age for each of the 96 states with an overall decreasing trend as the age at which the developmental rate curve first crosses or approaches zero. For the monotonically decreasing tasks, it is the age at which the developmental rate reaches its minimum absolute value, marking the beginning of a plateau. For tasks with transient recovery, the optimal age is the age of the first local minimum on the developmental curve, marking the end of the initial decline before the brief rebound.

**Fig. 3.**
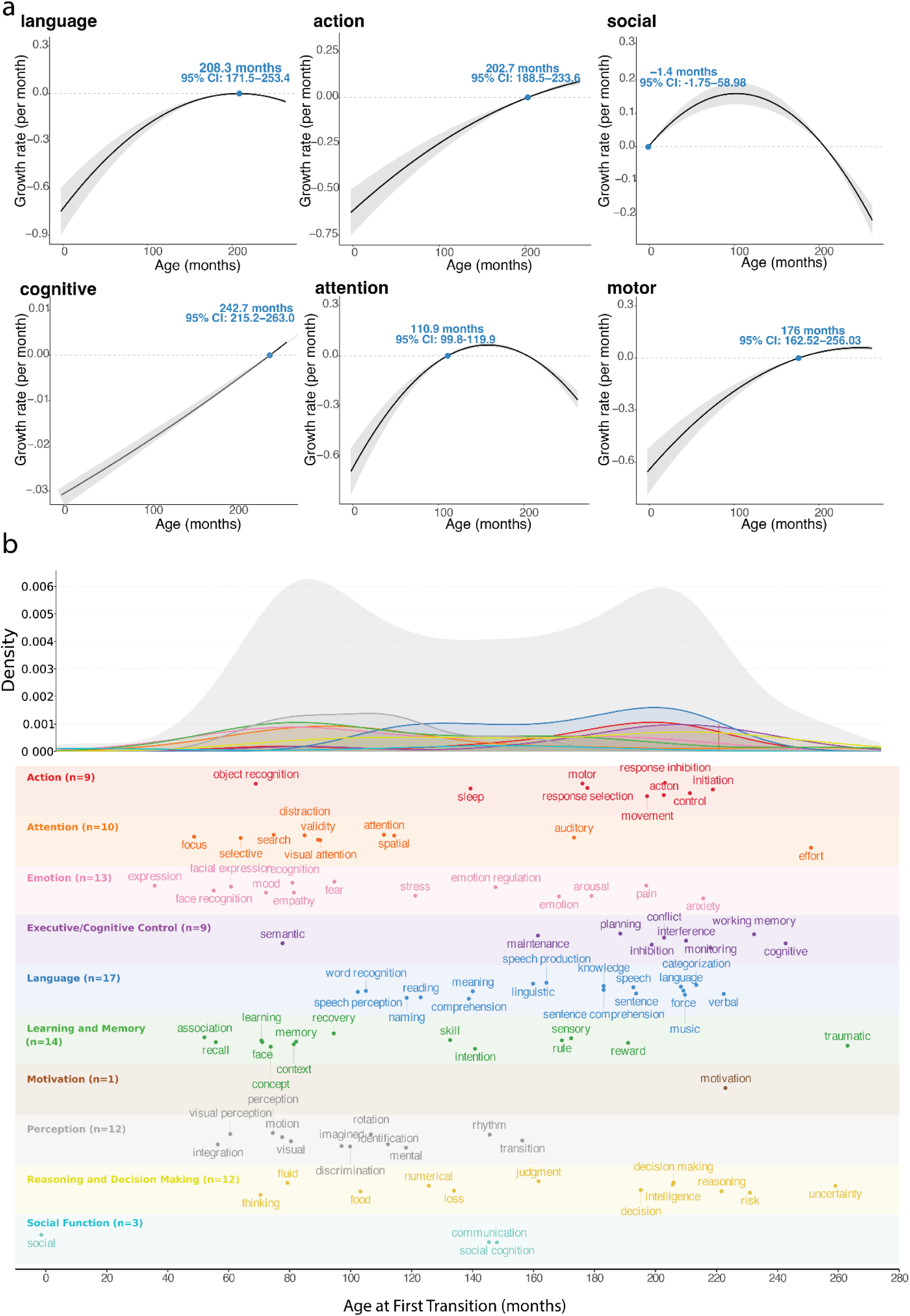
Optimal ages of cognitive energy efficiency and the distribution of first transitions. **a,** Growth rate curves (first derivative of the fitted GAMLSS trajectory) for six representative cognitive tasks. The blue dot in each panel marks the age at which the growth rate crosses zero. **b,** Kernel density estimate of the distribution of optimal ages across all cognitive tasks, color-coded by cognitive domain, with an early peak in childhood (∼35–120 months) and a late peak in adolescence (∼180–240 months).

As shown in Fig. 3b, optimal ages varied widely across the 96 tasks (range: −1.4 to 263.0 months; mean: 141.5 months; median: 140.4 months). The distribution was broadly bimodal, with one early peak concentrated in childhood (approximately 35–120 months) and a late peak in adolescence (approximately 180–240 months). Tasks related to perceptual and socio-affective processing, including face recognition, emotion, visual perception, and memory, reached their optimal age earlier (mean: 100.2 months), suggesting that these systems mature and stabilize relatively rapidly during childhood. In contrast, tasks associated with executive and regulatory functions, including cognitive control, working memory, inhibition, reasoning, and decision-making, reached their optimal age markedly later (mean: 205.5 months), consistent with the protracted maturation of the frontoparietal network extending into late adolescence and early adulthood. These findings suggest a hierarchical temporal organization of cognitive development, in which lower-order perceptual and social processes become energetically efficient well before higher-order executive systems.

To investigate how neurodevelopmental events shape the control energy landscape, we replaced the uniform control input **B**_*K*_ with region-specific weights derived from gene expression profiles of six neurodevelopmental processes ^33^: myelination, axonal development, dendrite development, synaptogenesis, neuron migration, and neuron differentiation (Fig. 4). Gene expression was obtained from the Allen Human Brain Atlas and parcellated into AAL2 regions using the abagen toolbox (v0.1.3)^34^. Statistical significance was assessed against 1,000 spin-test null models preserving spatial autocorrelation (Table S4).

**Fig. 4.**
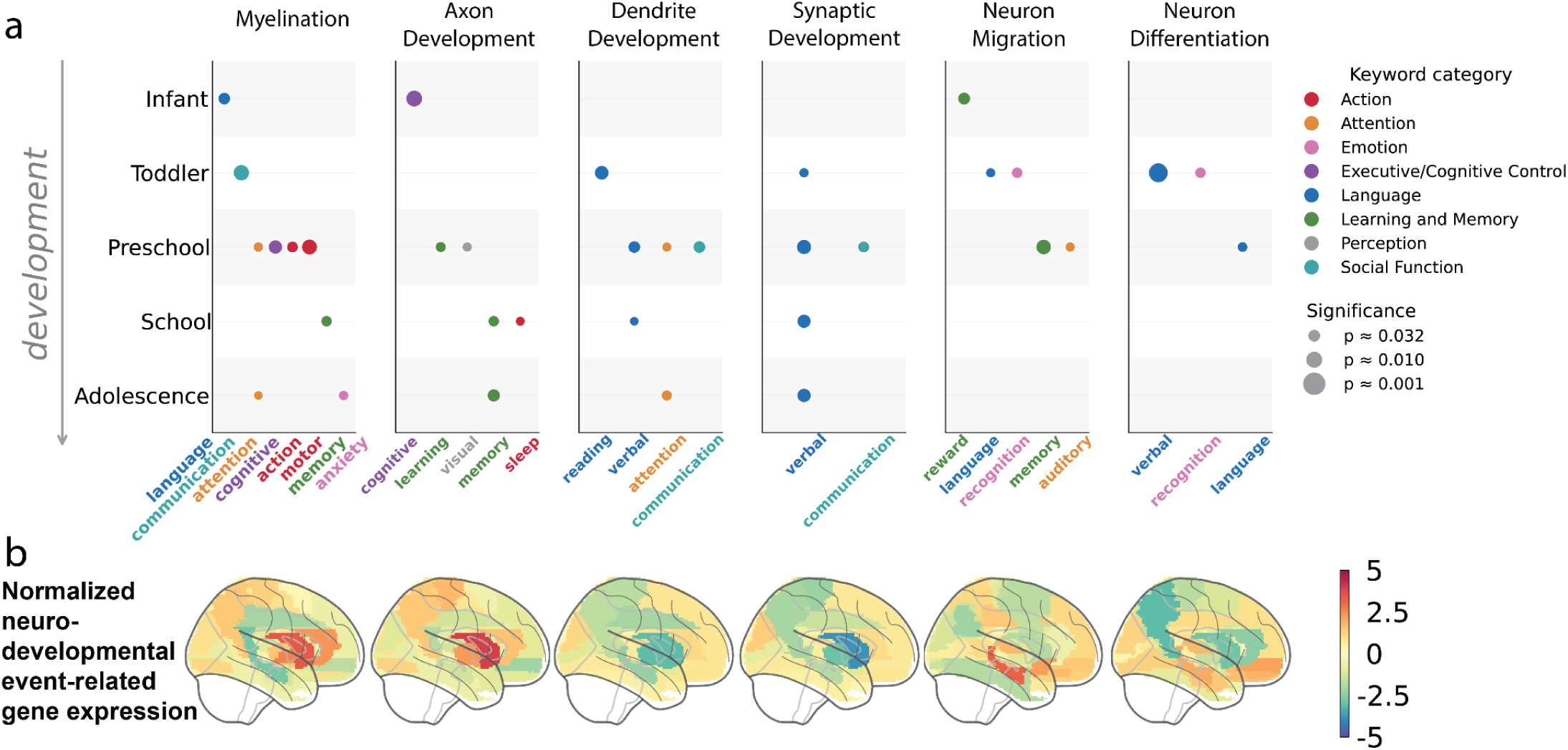
Neurodevelopmental events shape the control energy landscape on distinct biological timetables. **a,** Heatmap showing the significant influence of six neurodevelopmental gene expression profiles on the control energy for specific cognitive tasks across five developmental periods. Bar intensity indicates the magnitude of the effect as assessed by spin-test permutation against 1,000 spatially autocorrelated null models. Only tasks with significant effects (p_spin < 0.05) are displayed. **b,** Normalized gene expression maps for each of the six neurodevelopmental events, projected onto the cortical surface. Color scale indicates z-scored expression level (red = high, blue = low), illustrating the distinct spatial distribution of each event across the brain.

The six events differed remarkably in the temporal breadth and cognitive scope of their influence, following a pattern of temporal correspondence: events with earlier developmental onset and shorter duration exerted more restricted effects, whereas events with protracted time courses showed broader and more sustained influence.

Neuron migration and differentiation, both predominantly prenatal processes, affect the cognitive energy landscape within the most limited age range. Neuron migration significantly modulated control energy from infancy through the preschool period, influencing reward (p_spin = 0.018) in infancy; language (p_spin = 0.041), recognition (p_spin = 0.028), and memory (p_spin = 0.018) in the toddler period; and auditory processing (p_spin = 0.040) in the preschool period. No significant effects were found thereafter. Neuron differentiation exerted an even more restricted influence, with significant effects confined to the toddler period for recognition (p_spin = 0.029) and verbal processing (p_spin = 0.001), and to the preschool period for language (p_spin = 0.036). These patterns suggest that the spatial distribution of neurons established prenatally continues to constrain the postnatal energy landscape, but that this influence is progressively superseded by later maturational processes.

Axonal development’s strongest effect appeared in infancy for cognitive processing (p_spin = 0.004), with additional effects on learning (p_spin = 0.032) and visual processing (p_spin = 0.040) in the preschool period, sleep (p_spin = 0.044) and memory (p_spin = 0.029) at school age, and memory (p_spin = 0.016) in adolescence. Although spanning four of five developmental periods, the number of tasks affected per period was small (n=1–2), consistent with a focused but persistent contribution from long-range axonal connectivity.

Dendrite development and synaptogenesis, both involved in local circuit elaboration, showed effects beginning in toddlerhood and persisting into adolescence. Dendrite development influenced reading (p_spin = 0.010) in toddlerhood, attention (p_spin = 0.042) and communication (p_spin = 0.019) in the preschool period, verbal processing (p_spin = 0.048) at school age, and attention (p_spin = 0.031) in adolescence. Synaptogenesis showed a notably consistent influence on verbal and communicative functions: verbal processing in the toddler (p_spin = 0.040), preschool (p_spin = 0.008), school-age (p_spin = 0.012), and adolescent (p_spin = 0.011) periods, as well as communication in the preschool period (p_spin = 0.025). This domain specificity suggests that synaptic remodeling plays a particularly prominent role in shaping the neural efficiency of language-related circuits.

Myelination exhibited the most widespread and prolonged influence and was the only event with significant effects across all five developmental periods. Its effects spanned nine distinct cognitive tasks: language (p_spin = 0.020) in infancy; communication (p_spin = 0.005) in toddlerhood; motor (p_spin = 0.006), cognitive control (p_spin = 0.011), action (p_spin = 0.026), and attention (p_spin = 0.039) in the preschool period; memory (p_spin = 0.029) at school age; and anxiety (p_spin = 0.038) and attention (p_spin = 0.047) in adolescence. This prolonged, domain-general influence aligns with the long developmental time course of myelination, which starts from the prenatal period and extends into the third decade of life. The broad influence also underscores its unique role as a sustained mechanism for enhancing the efficiency of neural transmission across the cognitive landscape.

Finally, we estimated the optimal control energy required for each of the 10,000 possible transitions between pairs of cognitive states across all 3712 subjects. We then averaged the transition energy matrices across participants to characterize group-level patterns. Control energy varied substantially across transitions, spanning an approximately 17-fold range. Transitions toward the “emotional regulation” state required the most control energy, whereas transitions toward the “focus” state required the least control energy (Fig. 5a; Fig. S6 with each row and column labelled).

**Fig. 5.**
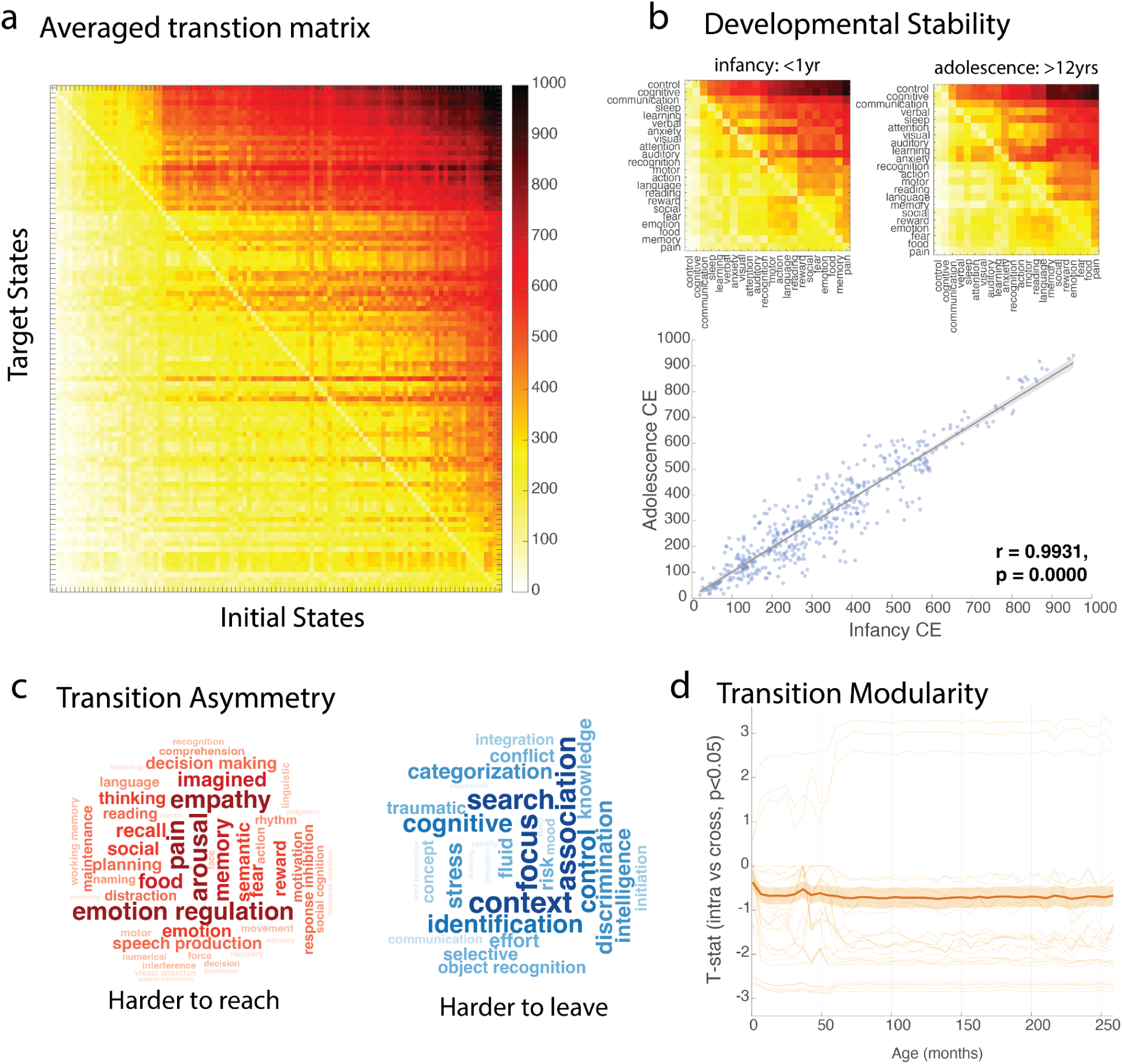
Control energy required to transition between brain cognitive states. **a,** Transition energy matrix showing the control energy cost of switching between all pairs of 100 cognitive states averaged across development. Rows represent initial states; columns represent target states. **b,** Developmental stability of the transition architecture. Transition matrices between cognitive states of high-frequency keywords are shown for infancy (<1 year) and adolescence (>12 years). The scatter plot below shows the correlation between infancy and adolescence control energy for all transition pairs (r = 0.933, p < 0.0001). **c,** Transition asymmetry, visualized as word clouds of cognitive states that are significantly harder to reach (left, red) versus harder to leave (right, blue). Word size reflects the magnitude of the asymmetry (t-statistic). **d,** Developmental trajectory of transition modularity. Each line represents a cognitive task, with the y-axis showing the t-statistic comparing within-category versus cross-category transition costs over age (x-axis, in months).

We observed transition asymmetry at two levels. First, at the landscape level, the target state played a more prominent role than the initial state in determining energy cost: values in the upper triangle of the control energy transition matrix were significantly higher than those in the lower triangle (t=9.36, p=1.20e-20), indicating systematic directional asymmetry. Second, at the state-specific level, we compared, for each cognitive state, the average energy required to reach it (energy to reach) versus the energy required to leave it (energy to leave; Fig 5c). Of the 100 cognitive states, 83 showed significant differences between energy-to-reach and energy-to-leave. Among these, 49 states were harder to reach, whereas 34 were harder to leave (Table S5). Notably, 11 of the hard-to-reach states belonged to the language category, whereas hard-to-leave states were distributed more broadly across cognitive domains. Similar transition patterns were observed across developmental periods, with extremely high correlations between any age groups and the overall average matrix (r > 0.99 for all comparisons; Fig. 5b; Fig. S7), indicating strong developmental stability in transition energy architecture.

To determine whether transitions are energetically more favorable within the same cognitive category defined by Cognitive Atlas, and how this property (task modularity) evolves across development, we compared the control energy required to transition from a given state to states within the same category versus states in different categories (two-sample t-tests) and used the t-statistics to represent the task modularity level (Fig. 5d). Across the entire developmental period, transitions originating from states such as attention, distraction, spatial processing, visual attention, and memory required significantly less energy when directed toward states within the same category, indicating a category-constrained transition architecture. Importantly, this within-category facilitation strengthened with age. During infancy, only five cognitive states exhibited significantly lower within-category transition costs, whereas by adolescence, more than 20 topographies demonstrated this pattern, suggesting increasing functional segregation and category-specific stabilization over development. Notably, three cognitive states—recognition, pain, and empathy—showed the opposite pattern, requiring less energy to transition to states outside their own category. This cross-category facilitation suggests that these functions may serve integrative or hub-like roles within the broader cognitive landscape (Fig. S8).

### Sensitivity analysis

To test whether the observed developmental pattern of control energy reduction is robust to the choice of meta-analytic database, we compared trajectories derived from NeuroSynth and BrainMap for the 11 cognitive states represented in both databases: action, attention, emotion, inhibition, language, memory, music, pain, perception, reasoning, and spatial. Among these 11 cognitive states, all showed converging evidence of a developmental decrease in control energy, indicating a robust pattern of control energy reduction. Mean optimal ages across the overlapped 11 cognitive states are similar (NeuroSynth: 16.4 years old and BrainMap: 16.2 years old), suggesting a consistent peak time of cognitive maturation during late adolescence. Notably, inhibition showed nearly identical inflection ages in NeuroSynth (16.6 years old) and BrainMap (16.7 years old). Similar results can be found in the cognitive tasks, including spatial, action, music, and reasoning (Fig. S9).

## Discussion

Leveraging network control theory and approximately 4,000 developmental diffusion MRI scans, we constructed a comprehensive energy landscape for 100 cognitive states from infancy through adolescence. The control energy to drive the majority of cognitive tasks decreased over development, each with a distinct temporal trajectory. Ages of optimal energy efficiency followed a bimodal distribution, peaking around school age and late adolescence, where social and perceptual functions reach efficiency earliest. Molecular-level neurodevelopmental events differed systematically in the temporal breadth and cognitive scope of their influence on the energy landscape. The cognitive transition architecture was remarkably stable across development yet showed progressive modularization.

Up to this point, a comprehensive growth chart of brain development supporting cognitive functions has remained elusive. This approach went beyond the limited task-based brain activity captured in each neuroimaging dataset and enabled exploration of the full range of possible brain activations within the scope of developmental cognitive science. Taken together, our findings provide an overall picture of how the cognitive energy landscape forms during development, reflecting the support of brain structural maturation for various cognitive functions in a timely order.

### Heterogeneous developmental trajectories reflect the temporal logic of cognitive demand

A central finding of this work is that the developmental trajectories of control energy were not uniform across cognitive tasks: while the majority decrease over development, they follow distinct temporal paths that cluster into five dissociable groups. This heterogeneity might reflect an adaptive temporal logic shaped by the ecological demands of each developmental period. Children do not learn all cognitive skills at once; an infant needs sensory discrimination and social attunement to locate their caregiver to survive rather than other higher-order cognitive functions^35,36^. The staggered reduction of control energy that we observe across domains may therefore represent a learning strategy of the human brain allocating its limited resources, from metabolic energy to myelination, to the brain regions and circuits most urgently needed at each stage of life^37,38^. Interestingly, we find that the trajectories can be steadily clustered, and the data-driven trajectory clusters did not neatly map onto canonical cognitive categories. This divergence may reflect the intertwined nature of cognitive development; for example, social skills can help boost children’s language performance^39,40^, and children with language challenges can find it more difficult to make friends and socialize^41,42^.

Notably, over half of the cognitive tasks exhibited a transient increase in control energy mid-trajectory, then resumed their decline. This temporary rise may reflect the energetic cost of active skill acquisition: before a cognitive ability is learned, the brain must recruit and coordinate circuits not yet optimized for the task, potentially demanding greater control energy than either the naive state that preceded it or the proficient state that will follow. Once the skill is consolidated through synaptic strengthening, pruning of inefficient connections^43^, and myelination of task-relevant pathways^44^, the energy cost declines below its original baseline. This learning-drive hypothesis parallels other U-shaped behavioral phenomena in developmental psychology, suggesting a transient cost of building a more powerful cognitive system^45^.

### The earliest cognitive energy efficiencies serve the social needs of infancy

We found that the cognitive functions reaching their optimal ages of brain energy efficiency earliest are not sensory or motor, as classical accounts of cortical maturation might predict, but social-related functions; the top five earliest-maturing tasks are heavily social-oriented, including social, expression, focus, association and facial recognition. This ordering aligns with the emerging evidence that humans are social beings from birth: neonates preferentially orient to face-like stimuli within hours of birth^35,46^, two-month-old infants engage in protoconversational turn-taking^47^, and by three months they can discriminate emotional expressions^48^. Even before birth, this distinct reaction to social signals has been recorded: fetuses increase their heart rates when hearing their mother’s voice at 32 weeks postmenstrual age^49^. These cognitive capacities are arguably the most basic needs for survival in a species as altricial as ours, which essentially depends on the infant’s ability to elicit and maintain caregiving by establishing social engagement with caregivers.

Our energy landscape results provide a brain-level account for this priority: the white-matter structure supporting social and face-related processing reaches an energy-efficient configuration earlier than that supporting other cognitive domains. This finding is consistent with the hypothesis that the infant brain is pre-organized, through a combination of genetic specification and prenatal experience, to prioritize the neural circuits most crucial to survival in the immediate postnatal environment^50,51^. The early efficiency of social circuits may also create a scaffold upon which later-developing capabilities, including language, theory of mind, and moral reasoning, are built^52^, consistent with developmental cascading models in which early social competence channels subsequent cognitive development^53^.

### Neurodevelopmental events shape the energy landscape in their own biological timetable

The transcriptional analysis revealed a systematic relationship between the time course of each molecular-level neurodevelopmental event and the temporal breath and cognitive scope of its influence on the cognitive energy landscape. This temporal correspondence provides new insight into how molecular-level processes translate into cognitive-level outcomes.

Neuron migration and differentiation, both largely completed before birth, exerted the most limited postnatal effects – confined to the toddler and preschool periods and affecting a narrow set of tasks^54^. Their restricted influence likely reflects how the prenatal spatial arrangement of neurons constrains brain network topology, but this constraint might be gradually overwritten by experience-driven neural plasticity^55^. On the other hand, myelination, which begins prenatally but lasts into the third decade of life, showed effects spanning all developmental periods and the widest range of cognitive domains^37^. This is consistent with the view that myelination acts as a domain-general mechanism for enhancing neural transmission efficiency and that its prolonged time course allows it to serve as a sustained driver of cognitive energy reduction across development^56,57^. Between these two extreme cases, synaptogenesis, dendritic and axonal development each contributed intermediate temporal layers. Notably, synaptogenesis selectively influenced only verbal and communicative functions across all developmental periods. This specificity may reflect the high dependence of language circuits on precise synaptic development^58^, which involves not only the initial overproduction of synapses but also experience-dependent pruning throughout childhood and adolescence^59^.

### A stable but increasingly modularized cognitive transition architecture

The transition energy matrix, capturing the energy cost of task switching between all pairs of cognitive states, revealed a mix of stability and change during development. The overall architecture of transition costs was remarkably consistent across developmental stages and similar to the results in adulthood^60^, indicating that the brain is born with a built-in map that determines which cognitive transitions are energetically more favorable and which are more costly. Yet, within this stable scaffold, we observed a constant increase in category-specific modularization. In infancy, five cognitive states showed significantly lower transition costs within their own cognitive categories; by adolescence, this number exceeded twenty. This developmental increase in within-category facilitation implies that experience and maturation progressively sharpen the boundaries between cognitive domains^61^, making it increasingly efficient to transition among related processes while maintaining higher barriers between unrelated ones. Evidence from functional connectivity studies has suggested increasing functional segregation during development^62^, which aligns with the results here in the structural connectome regarding control energy.

### Limitations

Several limitations should be noted when interpreting these results. First, the NeuroSynth activation maps used to define cognitive states are derived from fMRI studies regardless of the age ranges of the subjects^27^. Debates have been heard within the developmental cognitive science societies about whether children would follow adult-like brain activation patterns under certain tasks^9,63^. Future studies incorporating task-based fMRI collected directly in pediatric populations could help validate the developmental stability of these activation topographies. Second, the Allen Human Brain Atlas gene expression data collected from adult postmortem brains were used in the transcriptional analysis for their high spatial resolution. The regional expression profiles are therefore proxies for the developmental spatial distribution of neurodevelopmental gene sets, rather than direct measurements of gene expression at each age^64^. The assumption that relative spatial patterns of expression are conserved across development is supported by prior work^6,65^ but remains an approximation. Spatiotemporal gene expression atlases spanning prenatal through postnatal development, as they become available at sufficient spatial resolution, would enable more precise mapping of molecular influences onto the developmental energy landscape. Third, the linear discrete-time control model is a simplification of inherently nonlinear neural dynamics. While prior work has demonstrated that linear models capture a significant portion of variance in fMRI-derived brain dynamics^66^, the degree to which this approximation holds across developmental stages has not been systematically established, especially in the rapidly changing infant brain. Nonlinear extensions of network control theory may reveal dynamics not captured by the current framework. Finally, although the sample is large and multi-site, the data are predominantly cross-sectional, which limits inference about individual developmental trajectories. Longitudinal validation would strengthen the developmental claims and enable the detection of individual differences in the timing and shape of energy landscape maturation.

## Methods

### Neuroimaging Datasets

#### Participants

To outline the normative growth of control energy required to activate each cognitive state, we aggregated the available developmental neuroimaging datasets (i.e. dHCP^67^, BCP^68^, HCPD^69^, HBN^70^), each containing diffusion MRI scans for each participant and behavioral assessments tailored to the age range. For participants with more than one scan recorded, only the first scan was included in our analysis. Together, we included 3712 dMRI scans with 3455 participants, whose ages spanned from 27 postmenstrual weeks to 22 years. These scans were obtained from 11 sites in the four datasets. Participants demographics and imaging scan parameters for each site can be found in Supplementary Table 1 and 2. Written informed consent was obtained from participants and/or their legal guardians, and the recruitment procedures were approved by the local ethics committees for each dataset.

#### Image quality-control process

A standardized quality-control procedure is important to ensure the reliability and credibility of the neuroimaging data and, therefore, the growth curves of the control energy. Here, we implemented quality-control steps after every stage of diffusion MRI processing to construct the structural connectome: (1) quality control of raw images, (2) passage through the entire processing pipeline, and (3) neighboring DWI correction (NDC)^71^ after denoising. A total of 393 scans were excluded after all stages of screening, resulting in the final sample of 3,712 scans (Fig. S10).

#### Data processing pipeline

In order to unify the diffusion imaging process pipeline, we employed DSI-studio with the newly developed Fiber Data Hub to process the dMRI scans across the four datasets^72^. Despite efforts to ensure pipeline uniformity and comparability across the datasets, subtle processing differences were applied, especially for newborns in dHCP and infants in BCP. For newborns from dHCP, we applied the recommended diffusion SHARD pipeline to denoise and correct for Gibbs ringing, motion, eddy current, and susceptibility artifact^73^. For participants from BCP, artifacts were addressed by dmriprep, the recommended dMRI processing pipeline by BCP^74^. For individuals from HCPD and HBN, the susceptibility and eddy current artifacts were corrected using FSL topup and eddy^75^. After denoising and correction for artifacts, the image quality was checked using neighboring DWI correction (NDC)^71^. All diffusion data that passed NDC were reconstructed using generalized q-sampling imaging (GQI)^76^ and tensor metrics were calculated.

After reconstructing images with GQI, whole-brain fiber tracking was performed in DSI-studio (http://dsi-studio.labsolver.org/), using quantitative anisotropy (QA) as the termination index. QA values were computed in each voxel’s native space for every subject. The tracking parameters were set to an angular cutoff of 60 degrees, a step size of 1.0 mm, a minimum length of 30 mm, and a maximum length of 300 mm. The whole-brain fiber tracking process was performed with the FACT algorithm^77^ until 1,000,000 streamlines were reconstructed for each individual.

Age-matched versions of the AAL atlas were applied to construct the structural connectome for each participant. For dHCP participants, the neonatal version of the infant brain atlas was used^78^. For BCP participants, the neonatal, 1yr, or 2yr version of the infant brain atlas was used based on the age when the scans were performed^78^. For HBN and HCPD participants, the AAL2 atlas was used to construct the structural connectome^79^. T2-weighted images in native DWI space were used to inform region segmentation during connectome construction. The structural connectome for each individual was then constructed with a connectivity threshold of 0.001, and the pairwise connectivity strength was calculated as the average QA value of each fiber connecting the two end regions. Detailed DTI data preprocessing for each dataset is provided in the Supplementary Information.

### Defining cognitive states

Cognitive states of the association between voxels and cognitive keywords were obtained from NeuroSynth^27^, an automated term-based meta-analytic tool that synthesizes results from over 14000 published fMRI studies by searching for high-frequency keywords that are systematically mentioned in the papers alongside fMRI voxel coordinates (https://github.com/neurosynth/neurosynth, using the volumetric association test maps). This measure of association strength reflects the tendency for a given term to be reported in a functional neuroimaging study when activation is observed at a given voxel.

Although more than one thousand terms are cataloged in the NeuroSynth platform, we chose to focus on keywords in developmental cognitive science. To avoid manual selection bias, we implemented a two-step process: (1) find the NeuroSynth terms related to cognition, and (2) among those, find the terms most relevant to developmental cognitive science. For step 1, we created an overlapping set of the terms from NeuroSynth^27^ and the Cognitive Atlas^29^, a public ontology of cognitive science. This process resulted in 433 terms related to cognition. For step 2, we first created a text pool with all the questions in the 199 behavioral assessments collected in the dHCP, BCP, HBN, and HCPD datasets. We included repeated assessments to highlight joint interests in similar cognitive domains across different age ranges; for example, the Child Behavior Checklist (CBCL)^31^ was collected in every dataset and therefore, included four times in the whole set of assessments. Next, we counted the occurrences of 433 cognitive keywords in the overlapping set from step 1 and used the top 100 keywords that most frequently occurred in the developmental behavioral assessments as the keywords we investigated in this work.

With the developmental cognitive keywords, we then generated the activation map of each keyword using NeuroSynth, which can be interpreted as a quantitative representation of how regional fluctuations in activity are related to cognitive processes. Note that the activation maps generated by NeuroSynth do not distinguish between areas that are activated or deactivated in relation to the term of interest, nor the degree of activation, only that certain brain areas are frequently reported in conjunction with certain words.

To assess the robustness of the developmental control energy trajectories to the choice of meta-analytic database, we replicated the primary analysis using activation maps derived from BrainMap as target states (xf), in place of NeuroSynth association-test z-maps. This approach mirrors the sensitivity analysis reported by Luppi *et al.* 2024^60^. BrainMap activation maps were obtained by querying the BrainMap functional database^80,81^ using Sleuth 3.0.4. Searches were restricted to experiments with Context = Normal Mapping and Activation = Activations Only, using Behavioral Domain filters covering 11 cognitive states that overlap with the states defined with NeuroSynth: action, attention, emotion, inhibition, language, memory, music, pain, perception, reasoning, and spatial. For each state, exported coordinates (GingerALE format) were converted to NiMARE Datasets and submitted to Activation Likelihood Estimation (ALE) meta-analysis using NiMARE 0.0.13^82^, with sample-size-dependent FWHM kernels.

Unthresholded ALE z-maps were retained as activation maps, consistent with the unthresholded association-test z-maps produced by NeuroSynth, ensuring methodological comparability between databases. Regional activation vectors were extracted using the AAL2 atlas with cortical activations. Activation map vectors were binarized (threshold > 0) to define the target cognitive state xf, and the optimal control energy was computed for each subject using NCT parameters identical to those used in the primary analysis. The same downstream analyses were performed to validate the main results.

### Network control theory

The brain can be considered as a naturally occurring complex system^83^. One way to understand the behavior of a complex system is to understand the mechanisms that control it, which involves driving the system to the desired state. Network control theory was developed to determine how to control a system consisting of nodes (e.g., brain regions) and edges (e.g., white matter tracts between brain regions). The approach can be used to characterize both dynamic changes in the short term and developmental trajectories in the long term, in addition to distinguishing between cognitive dysfunctions in clinical groups^11^.

A network system can be represented as a graph **G** = (**V**, **E**), where **V** and **E** are the vertex and edge sets, respectively. Let a_ij_ be the weight associated with the edge (*i*, *j*) in **E** and define the weighted adjacency matrix of the graph **G** as **A** = [a_*ij*_], where *a*_*ij*_ = 0 when *a*_*ij*_ ∉ **E**. Here, the individual structural connectome **A** inℝ^*n*×*n*^ is a symmetric and weighted adjacency matrix whose elements [*a*_*ij*_] evaluate the strength of the white matter fiber connecting region *i* to region *j* in the brain.

For the definition of the neural dynamic processes, we adopt the prior models that link brain structural networks to simplified brain dynamics. Although the evolution of brain activity occurs in a nonlinear manner, previous studies have demonstrated that simplified linear models can predict a significant portion of the variance in neural dynamics recorded by fMRI^66^. Therefore, we employ a linear, time-invariant network model: 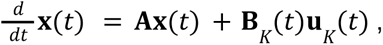 where ***x*** denotes the brain state at a given time, and **A** is the symmetric, undirected, and weighted adjacency matrix for the network. In our case, **A** represents the structural connectome for each individual, whose element indicates the pairwise strength of the structural connection. The diagonal elements are set to zero. The input matrix **B**_*K*_ identifies the control points *K* in the brain, where *K* = {*k*_1_, ···, *k*_*m*_} and **B**_*K*_ = [*e*_*k*1_··· *e*_*km*_]. *e* denotes the *i*th canonical vector of dimension *N*, and input ***u***_*K*_ denotes the input control strategy over time.

### Control energy to drive brain state transitions

To explore how the brain’s dynamic processes are constrained by the structural connectome, we utilized network control theory to model the energy required to activate each cognitive state defined as the meta-analytic activation map with regions associated with each cognitive keyword in the NeuroSynth database. In this case, the baseline state **x**(0) was set to zero and the target state **x**(*T*) was defined as a binary vector where all regions in the activation map had a magnitude of one, while all other regions had a magnitude of zero. We also investigated the energy required to transition between cognitive states. In this scenario, the only difference is the baseline state **x**(0) was set to the activation map of the starting state, and **x**(*T*) reflected the activation map of the target state.

Here, we followed the definition of the control task in the previous studies ^84,85^, where the system transitions from initial state to target state with minimum-energy input as an optimal control problem, with the cost function defined as a Hamiltonian, *H*(**p**, **x**, **u**, *t*) = **x**^T^**x** + **u**^T^**u** + **p**^T^(**Ax** + **Bu**). By solving the optimization problem **u**\* = *arg min*(*H*), control energy for each node *k*_*i*_ was defined as 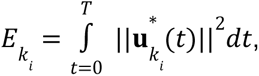 indicating the overall energy input required by the node to facilitate the desired state transition.

### GAMLSS model

To outline the trajectories of the control energy for developmental cognitive states, we used the GAMLSS model^86,87^ using the gamlss package (version 5.4-3) in R 4.2.0. Control energy for each cognitive state was modeled separately, with the control energy averaged across the whole brain as the dependent variable, age as a smoothing term using the B-spline basis function (degrees of freedom = 3), sex and the diffusion MRI quality-check value NDC as the fixed effects, and scanner sites as random effects.

To select an appropriate distributional family, we evaluated ten candidate distributions — Normal (NO), Student’s t (TF), Power Exponential (PE), Log-Normal (LOGNO), Gamma (GA), Inverse Gaussian (IG), Box-Cox t (BCT), Box-Cox Power Exponential (BCPE), Johnson’s SU (JSU), and Skew t type 3 (ST3) — fitted to each of the 100 outcomes. All ten families converged for all outcomes. Model fit was assessed using the Akaike Information Criterion (AIC); BCT achieved the lowest AIC in 42 outcomes and was within ΔAIC ≤ 2 of the best-fitting family in 62 outcomes, making it the most consistently well-fitting family overall (Fig. S11). To ensure comparability of distributional parameters across outcomes, BCT was adopted as a uniform family for all 100 models. This choice is also aligned with the World Health Organization guidelines for modeling anthropometric growth charts^87^.

The BCT distribution has four parameters: median (μ), coefficient of variation (σ), skewness (ν), and kurtosis (τ). The control energy for each cognitive state, denoted by y, was then modeled according to:

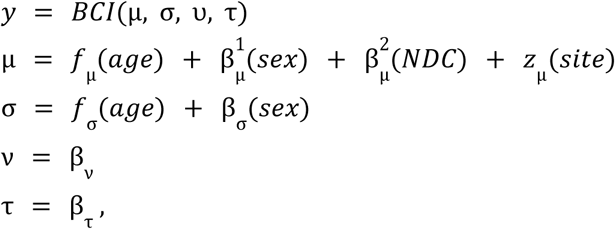

where *f*_μ_ and *f*_σ_ denote cubic B-spline smoothing functions of age (degrees of freedom = 3), β terms denote fixed-effect coefficients, and *z*_µ_ (*site*) denotes the scanner site modeled as a random effect on µ. Skewness (ν) and kurtosis (τ) were modeled with intercept-only terms, following the practice of previous lifespan neuroimaging studies^88,89^. For model estimation, the default convergence criterion (a log-likelihood difference of 0.001 between iterations) was used, with a maximum of 100 iterations. All 100 models converged successfully.

The developmental rate of each control energy trajectory was assessed by numerically differentiating the 50th centile curve, using a central difference approximation with a step size of 1e-5. Optimal ages were defined as the ages at which the growth rate crossed zero or was closest to zero if the trajectory decreased monotonically. To quantify uncertainty around these optimal ages, we performed bootstrap resampling (200 resamples), refitting the full GAMLSS model on each resample and recomputing the zero-crossing age. The 95% confidence interval (CI) for each optimal age was derived from the 2.5^th^ and 97.5^th^ percentiles of the bootstrap distribution. The goodness of fit of each normative model was evaluated using multiple complementary criteria. The Akaike Information Criterion (AIC) was used to assess distributional fit, and out-of-sample metrics including the coefficient of determination (R²), mean absolute error (MAE), and root mean squared error (RMSE) were used to evaluate predictive accuracy. Model adequacy was further assessed using the Shapiro–Wilk test applied to the normalized quantile residuals of each fitted model; all 100 models yielded W > 0.99, indicating that the residuals closely followed a normal distribution and confirming good distributional fits across all cognitive state outcomes. These metrics are reported for all 100 node energy outcomes in the Table S2.

### Transcriptional analysis

Gene expression levels were defined using the Allen Human Brain Atlas (AHBA)^64^. After preprocessing with the abagen (v0.1.3) toolbox^34^, we generated a gene expression matrix with rows as brain regions of the AAL2 atlas and columns as gene-expression levels. Six gene sets were selected following Kang *et al.* (2011) to characterize the following neurodevelopmental events: axonal development, dendrite development, neuron migration, neuron differentiation, myelination, and synaptogenesis^33^. To assess statistical significance, 1,000 null models were constructed for each event using the spin test that randomly rotated the original neurodevelopmental brain map while preserving its mean, variance, and spatial autocorrelation^90,91^. Significance was then determined by comparing the control energy calculated from the original gene expression map against that of the 1,000 null permutations.

## Data availability

The developmental neuroimaging datasets are available upon request from the developing Human Connectome Project (https://www.developingconnectome.org/data-release/), the Baby Connectome Project (https://nda.nih.gov/), the Lifespan Human Connectome Project (https://nda.nih.gov/), and the Healthy Brain Network (https://fcon_1000.projects.nitrc.org/indi/cmi_healthy_brain_network/).

## Code availability

Customized code in this study can be found in https://github.com/huiliii/cognitive_energy_landscape. Network control theory code can be found at https://nctpy.readthedocs.io/en/latest/. DSI-studio processing code can be found at https://dsi-studio.labsolver.org/. R code for gamlss modeling can be found at https://www.gamlss.com/.

## Acknowledgement

This work was supported by the ARO award W911NF-24-1-0228, NIH R01MH126133 and NIH R01MH137609.

## Notes

### Competing Interest Statement

The authors have declared no competing interest.

