## Supplementary Information for "Development of the cognitive energy landscape from infancy to adolescence"

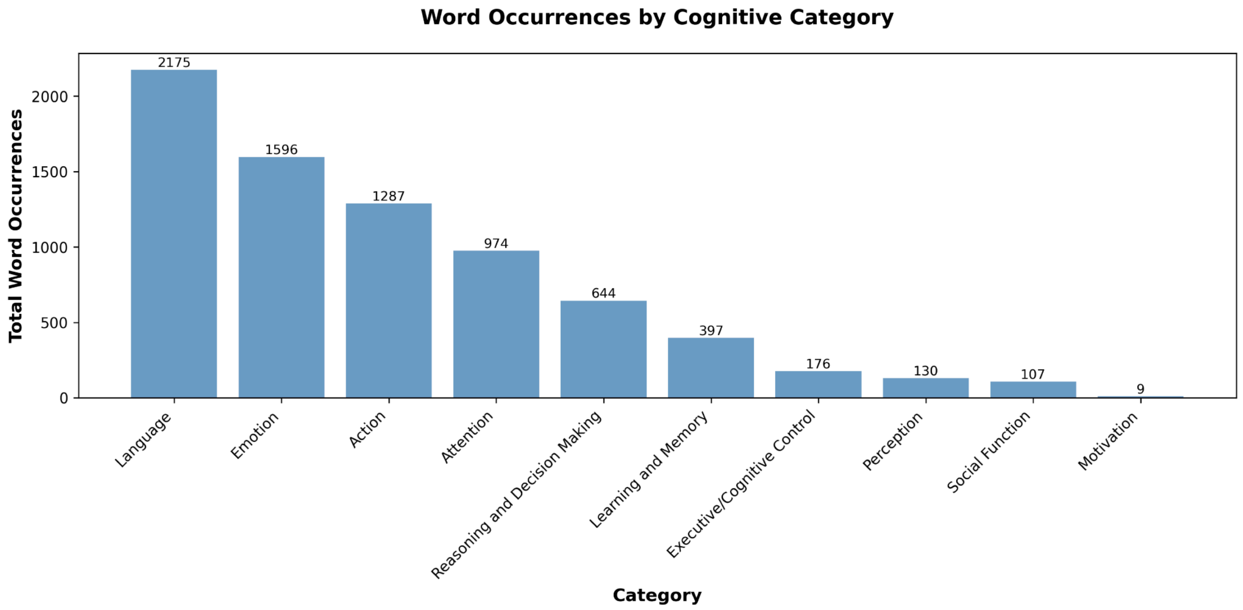


**Fig. S1. Cognitive keyword occurrences for each category in the Cognitive Atlas.** Each bar represents the total number of cognitive keyword occurrences in each Cognitive Atlas category. Language-related terms were the most frequent (n = 2,175), followed by emotion (n = 1,596), action (n = 1,287), and attention (n = 974).

**Table S1. Top 100 cognitive keywords and their categories defined in the Cognitive Atlas sorted by their occurrence**

| Keyword | Category | Keyword | Category |
|---|---|---|---|
| attention | Attention | fear | Emotion |
| skill | Learning and Memory | context | Learning and Memory |
| identification | Perception | action | Action |
| visual perception | Perception | empathy | Emotion |
| facial expression | Emotion | force | Language |
| uncertainty | Reasoning and Decision Making | concept | Learning and Memory |
| visual | Perception | imagined | Perception |
| speech | Language | discrimination | Perception |
| initiation | Action | spatial | Attention |
| numerical | Reasoning and Decision Making | control | Action |
| sleep | Action | mood | Emotion |
| face recognition | Emotion | movement | Action |
| reward | Learning and Memory | maintenance | Executive/Cognitive Control |
| linguistic | Language | recovery | Learning and Memory |
| recall | Learning and Memory | effort | Attention |
| traumatic | Learning and Memory | rhythm | Perception |
| reasoning | Reasoning and Decision Making | fluid | Reasoning and Decision Making |
| loss | Reasoning and Decision Making | sentence | Language |
| response inhibition | Action | focus | Attention |
| verbal | Language | social | Reasoning and Decision Making |
| intention | Learning and Memory | emotion | Emotion |
| motor | Action | search | Attention |
| association | Learning and Memory | music | Language |
| intelligence | Reasoning and Decision Making | selective | Attention |
| word recognition | Language | integration | Perception |
| perception | Perception | decision | Reasoning and Decision Making |
| knowledge | Language | food | Reasoning and Decision Making |

| | | | |
|---|---|---|---|
| judgment | Reasoning and Decision Making | sentence comprehension | Language |
| distraction | Attention | monitoring | Executive/Cognitive Control |
| thinking | Reasoning and Decision Making | language | Language |
| categorization | Language | working memory | Executive/Cognitive Control |
| recognition | Emotion | rotation | Perception |
| social cognition | Social Function | conflict | Executive/Cognitive Control |
| object recognition | Action | arousal | Emotion |
| expression | Emotion | motivation | Motivation |
| pain | Emotion | face | Learning and Memory |
| response selection | Action | comprehension | Language |
| speech perception | Language | mental | Perception |
| interference | Executive/Cognitive Control | rule | Learning and Memory |
| speech production | Language | visual attention | Attention |
| cognitive | Executive/Cognitive Control | stress | Emotion |
| naming | Language | semantic | Executive/Cognitive Control |
| sensory | Learning and Memory | anxiety | Emotion |
| motion | Perception | learning | Learning and Memory |
| auditory | Attention | reading | Language |
| meaning | Language | risk | Reasoning and Decision Making |
| decision making | Reasoning and Decision Making | communication | Social Function |
| transition | Perception | memory | Learning and Memory |
| emotion regulation | Emotion | planning | Executive/Cognitive Control |
| validity | Attention | inhibition | Executive/Cognitive Control |

**Fig. S2 Developmental trajectories and growth rate curves of control energy for cognitive states of high-frequency keywords defined by NeuroSynth**

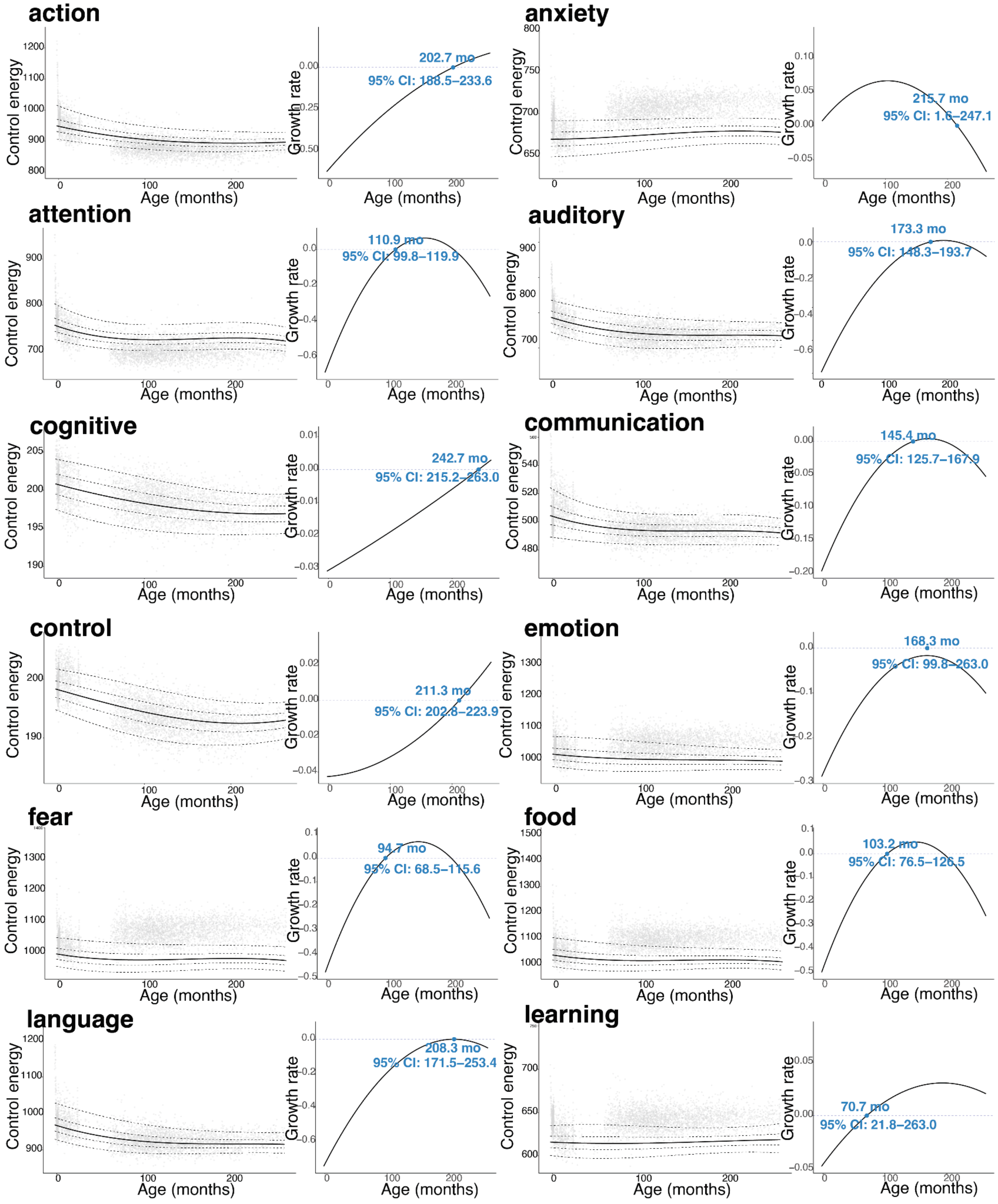

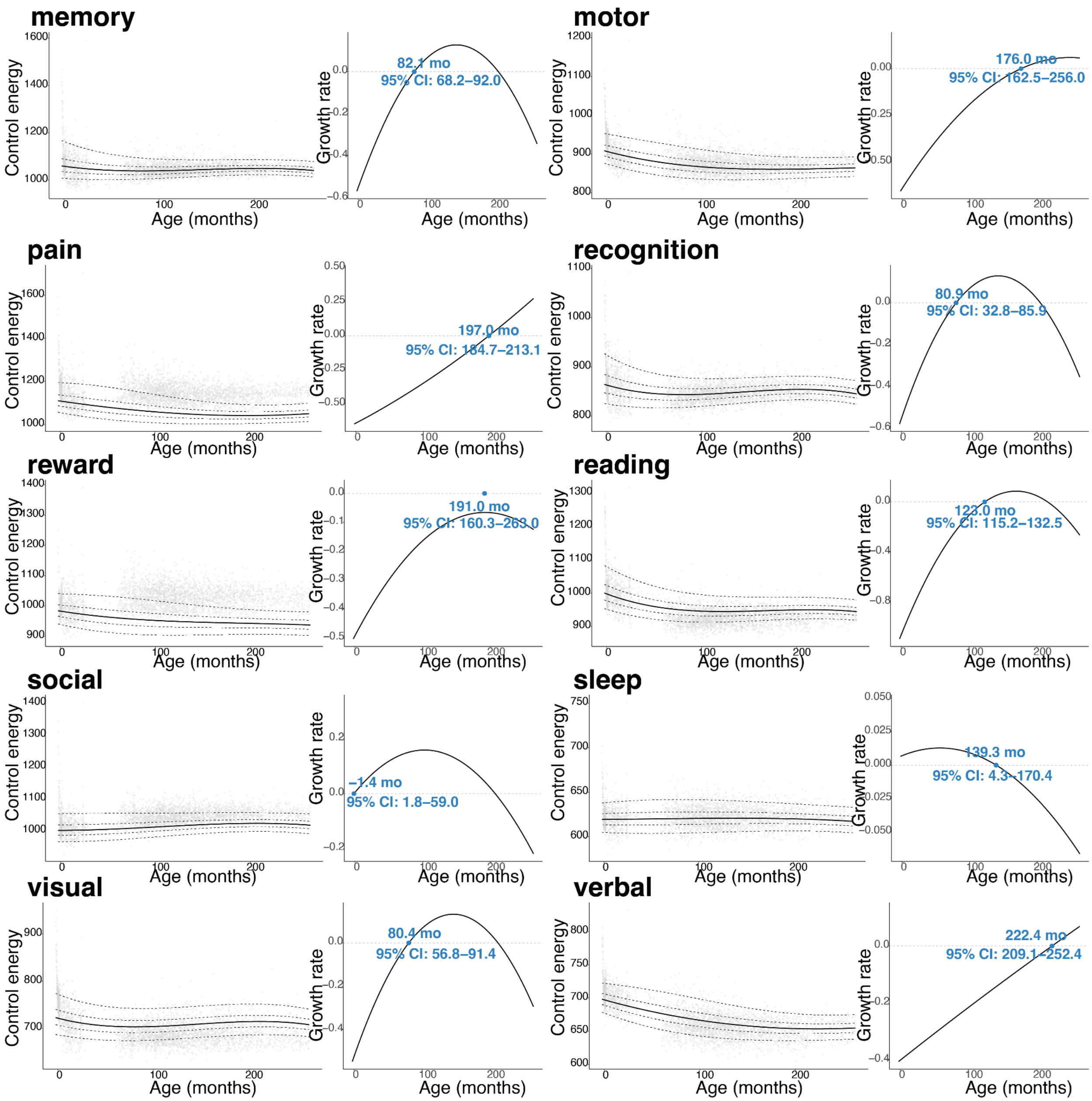
memory
Control energy
Age (months)
Growth rate
82.1 mo
95% CI: 68.2–92.0
motor
176.0 mo
95% CI: 162.5–256.0
pain
197.0 mo
95% CI: 184.7–213.1
recognition
80.9 mo
95% CI: 32.8–85.9
reward
191.0 mo
95% CI: 160.3–263.0
reading
123.0 mo
95% CI: 115.2–132.5
social
−1.4 mo
95% CI: 1.8–59.0
sleep
139.3 mo
95% CI: 4.3–170.4
visual
80.4 mo
95% CI: 56.8–91.4
verbal
222.4 mo
95% CI: 209.1–252.4

**Table S2. GAMLSS fitting performance for the developmental trajectories of the control energy of the 100 cognitive tasks**

| keywords | AIC | MAE | RMSE | Rsquared | Converged | keywords | AIC | MAE | RMSE | Rsquared | Converged |
|---|---|---|---|---|---|---|---|---|---|---|---|
| action | 35635.97 | 10.70 | 12.72 | 0.90 | TRUE | learning | 29746.01 | 6.15 | 7.43 | 0.48 | TRUE |
| control | 17978.88 | 1.87 | 2.39 | 0.64 | TRUE | linguistic | 34244.56 | 9.07 | 10.98 | 0.90 | TRUE |
| initiation | 25957.39 | 4.64 | 5.77 | 0.65 | TRUE | meaning | 33165.31 | 8.31 | 10.08 | 0.80 | TRUE |
| motor | 34886.93 | 9.74 | 11.66 | 0.81 | TRUE | music | 33338.97 | 8.56 | 10.36 | 0.55 | TRUE |
| movement | 35299.81 | 10.50 | 12.47 | 0.86 | TRUE | naming | 35065.66 | 9.81 | 11.69 | 0.76 | TRUE |
| object.recognition | 27247.19 | 5.73 | 7.16 | 0.52 | TRUE | reading | 36310.67 | 11.05 | 13.18 | 0.86 | TRUE |
| response.inhibition | 36465.70 | 11.59 | 13.71 | 0.79 | TRUE | sentence | 34031.08 | 9.14 | 11.01 | 0.67 | TRUE |
| response.selection | 32950.87 | 7.91 | 9.67 | 0.79 | TRUE | sentence.comprehension | 31593.54 | 7.77 | 9.48 | 0.71 | TRUE |
| sleep | 29375.78 | 6.11 | 7.41 | 0.18 | TRUE | speech | 33862.83 | 9.74 | 11.65 | 0.70 | TRUE |
| anxiety | 30412.71 | 6.55 | 7.84 | 0.73 | TRUE | speech.perception | 32506.62 | 7.75 | 9.36 | 0.71 | TRUE |
| arousal | 38643.71 | 12.61 | 14.85 | 0.60 | TRUE | speech.production | 36401.52 | 10.79 | 12.90 | 0.82 | TRUE |
| emotion | 37025.65 | 11.28 | 13.45 | 0.58 | TRUE | verbal | 31976.52 | 8.07 | 9.80 | 0.85 | TRUE |
| emotion.regulation | 39017.19 | 12.93 | 15.35 | 0.73 | TRUE | word.recognition | 30536.10 | 7.10 | 8.50 | 0.74 | TRUE |
| empathy | 37053.14 | 10.11 | 12.06 | 0.59 | TRUE | cognitive | 16385.31 | 1.56 | 1.97 | 0.34 | TRUE |
| expression | 30167.81 | 6.56 | 7.94 | 0.45 | TRUE | conflict | 26615.65 | 5.32 | 6.84 | 0.60 | TRUE |
| face.recognition | 31656.46 | 7.51 | 9.10 | 0.55 | TRUE | inhibition | 31299.65 | 7.35 | 8.86 | 0.65 | TRUE |
| facial.expression | 32040.42 | 8.15 | 9.85 | 0.40 | TRUE | interference | 34305.39 | 9.49 | 11.25 | 0.83 | TRUE |
| fear | 37327.01 | 11.03 | 13.12 | 0.76 | TRUE | maintenance | 34509.02 | 8.48 | 10.10 | 0.84 | TRUE |
| mood | 27109.12 | 5.15 | 6.35 | 0.56 | TRUE | monitoring | 32289.38 | 7.85 | 9.57 | 0.74 | TRUE |
| pain | 39607.75 | 14.71 | 17.47 | 0.61 | TRUE | planning | 36283.16 | 10.69 | 12.68 | 0.91 | TRUE |
| recognition | 33966.15 | 8.93 | 10.74 | 0.54 | TRUE | semantic | 35484.50 | 8.98 | 10.83 | 0.50 | TRUE |
| stress | 18111.74 | 1.91 | 2.47 | 0.29 | TRUE | working.memory | 34852.60 | 9.46 | 11.29 | 0.91 | TRUE |
| association | 9799.99 | 0.67 | 0.86 | 0.17 | TRUE | communication | 26732.75 | 5.00 | 6.22 | 0.53 | TRUE |
| concept | 24471.90 | 4.02 | 5.08 | 0.36 | TRUE | social.cognition | 33770.84 | 8.16 | 10.25 | 0.35 | TRUE |
| context | 6664.18 | 0.47 | 0.58 | 0.21 | TRUE | decision | 36164.51 | 11.90 | 14.15 | 0.39 | TRUE |
| face | 35184.22 | 9.72 | 11.77 | 0.42 | TRUE | decision.making | 37274.15 | 11.81 | 14.00 | 0.50 | TRUE |
| intention | 29336.76 | 6.25 | 7.81 | 0.34 | TRUE | fluid | 22617.37 | 3.39 | 4.28 | 0.35 | TRUE |
| recall | 35548.23 | 8.99 | 10.79 | 0.35 | TRUE | food | 37794.96 | 12.08 | 14.25 | 0.73 | TRUE |
| recovery | 33048.12 | 8.47 | 10.37 | 0.44 | TRUE | intelligence | 22006.04 | 3.13 | 3.95 | 0.23 | TRUE |
| reward | 38425.92 | 13.55 | 16.08 | 0.61 | TRUE | judgment | 33369.64 | 8.64 | 10.33 | 0.79 | TRUE |
| rule | 32915.86 | 8.36 | 10.15 | 0.73 | TRUE | loss | 33632.37 | 9.91 | 12.47 | 0.53 | TRUE |
| sensory | 33241.91 | 8.52 | 10.08 | 0.84 | TRUE | numerical | 34596.63 | 9.19 | 11.05 | 0.84 | TRUE |
| skill | 32604.99 | 8.44 | 10.11 | 0.74 | TRUE | reasoning | 32625.41 | 7.76 | 9.29 | 0.86 | TRUE |
| traumatic | 22482.86 | 3.17 | 3.97 | 0.18 | TRUE | risk | 23091.45 | 3.52 | 4.45 | 0.12 | TRUE |
| attention | 33641.82 | 9.32 | 11.20 | 0.86 | TRUE | social | 35484.42 | 9.09 | 10.83 | 0.74 | TRUE |
| auditory | 30951.90 | 7.21 | 8.66 | 0.77 | TRUE | thinking | 35348.90 | 9.34 | 11.17 | 0.20 | TRUE |

| | | | | | | | | | | | |
|---|---|---|---|---|---|---|---|---|---|---|---|
| distraction | 34554.23 | 8.63 | 10.36 | 0.54 | TRUE | uncertainty | 30573.09 | 6.81 | 8.19 | 0.40 | TRUE |
| effort | 23322.06 | 3.58 | 4.52 | 0.67 | TRUE | discrimination | 19238.47 | 2.19 | 2.82 | 0.52 | TRUE |
| focus | 5716.60 | 0.45 | 0.60 | 0.00 | TRUE | identification | 14457.71 | 1.22 | 1.56 | 0.10 | TRUE |
| memory | 36881.32 | 10.69 | 12.86 | 0.46 | TRUE | imagined | 36178.05 | 9.81 | 11.64 | 0.56 | TRUE |
| search | 13125.50 | 1.03 | 1.30 | 0.11 | TRUE | integration | 25548.98 | 4.49 | 5.58 | 0.45 | TRUE |
| selective | 27486.74 | 5.92 | 7.46 | 0.61 | TRUE | mental | 32939.19 | 8.32 | 10.12 | 0.79 | TRUE |
| spatial | 32466.00 | 8.57 | 10.37 | 0.77 | TRUE | motion | 33886.33 | 10.13 | 12.17 | 0.82 | TRUE |
| validity | 29500.51 | 6.21 | 7.54 | 0.27 | TRUE | perception | 30630.06 | 6.98 | 8.38 | 0.67 | TRUE |
| visual.attention | 35209.40 | 10.20 | 12.09 | 0.88 | TRUE | rhythm | 35134.11 | 9.73 | 11.57 | 0.79 | TRUE |
| categorization | 19516.63 | 2.25 | 2.84 | 0.33 | TRUE | rotation | 33668.48 | 9.47 | 11.39 | 0.86 | TRUE |
| comprehension | 33872.31 | 8.55 | 10.15 | 0.65 | TRUE | transition | 31352.77 | 6.91 | 8.28 | 0.80 | TRUE |
| force | 34241.82 | 9.27 | 10.99 | 0.79 | TRUE | visual | 34744.47 | 10.83 | 12.96 | 0.84 | TRUE |
| knowledge | 22426.70 | 3.16 | 4.03 | 0.28 | TRUE | visual.perception | 32901.56 | 8.68 | 10.41 | 0.60 | TRUE |
| language | 35665.10 | 10.37 | 12.63 | 0.79 | TRUE | motivation | 37703.35 | 13.41 | 16.09 | 0.38 | TRUE |

*Note: AIC = Akaike Information Criterion; MAE = mean absolute error; RMSE = root mean square error (both vs. fitted μ).*

**Table S3. K-means clustering results of the 100 cognitive tasks**

| | | | | | | |
|---|---|---|---|---|---|---|
| **Cluster 0** | Sleep | Response selection | Validity | Visual attention | Face recognition | Face |
| | Emotion | Linguistic | Verbal | Speech production | Sentence | Sentence comprehension |
| | Reward | Concept | Mood | Motivation | Loss | Judgment |
| | Thinking | Social | Decision | | | |
| **Cluster 1** | Response inhibition | Object recognition | Effort | Emotion regulation | Arousal | Uncertainty |
| | Monitoring | Speech perception | Naming | Meaning | Music | Identification |
| | Food | Intelligence | | | | |
| **Cluster 2** | Initiation | Movement | Focus | Expression | Pain | Communication |
| | Anxiety | Maintenance | Working memory | Planning | Knowledge | Categorization |
| | Memory | Learning | Sensory | Traumatic | Recall | Comprehension |
| | Motion | Fear | Rhythm | Reasoning | Risk | Discrimination |
| | Numerical | | | | | |
| **Cluster 3** | Action | Control | Attention | Distraction | Auditory | Recognition |
| | Search | Selective | Spatial | Empathy | Conflict | Facial expression |
| | Semantic | Inhibition | Speech | Force | Language | Reading |
| | Intention | Context | Visual attention | Visual | Perception | Transition |
| | imagined | Integration | Rotation | Decision making | Fluid | Social cognition |
| **Cluster 4** | Motor | Interference | Cognitive | Word recognition | Skill | Association |
| | Recovery | Rule | Mental | | | |

**Fig. S3 Results of the elbow method to determine the cluster number for the k-means clustering method.**

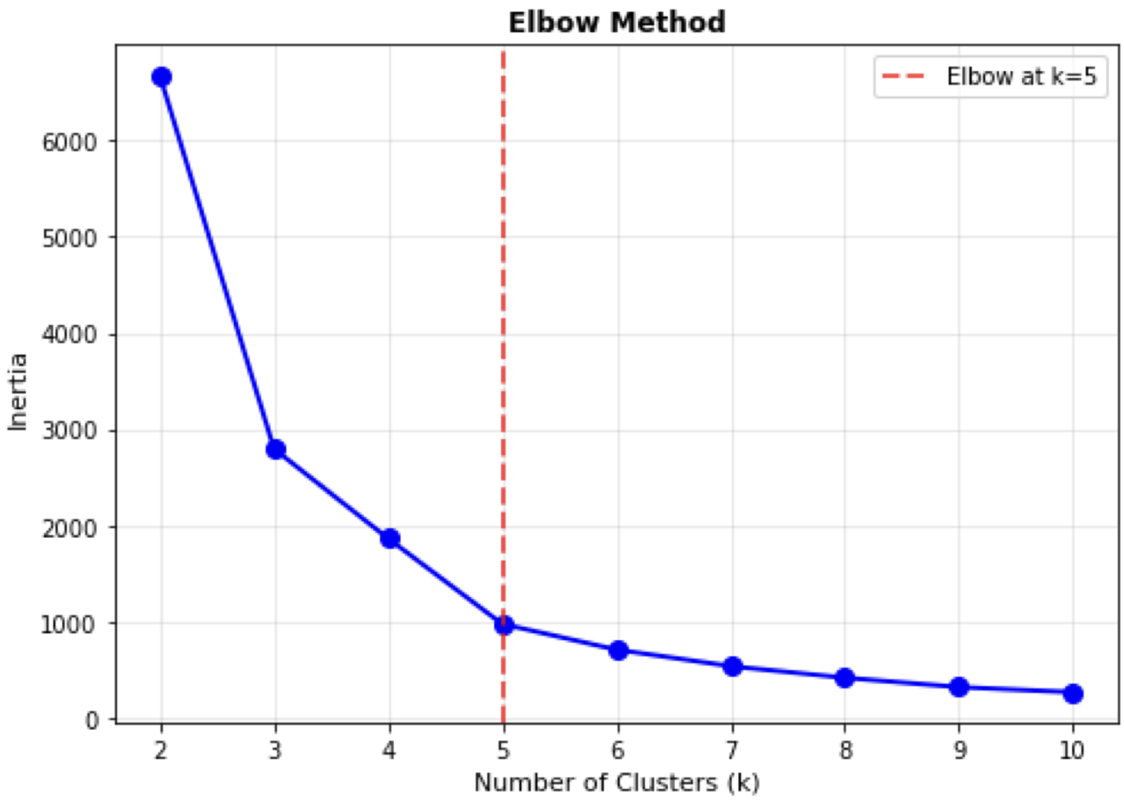


To identify the optimal number of clusters for the ke-mean clustering of the 100 control energy trajectories, the within-cluster inertia was computed across k = 2-10. The inertia curve exhibited the elbow point at k = 5 (red dashed line), supporting the selection of five clusters in the main analysis.

**Fig. S4. Pairwise agreement between clustering results using alternative clustering methods with the Adjusted Rand Index**

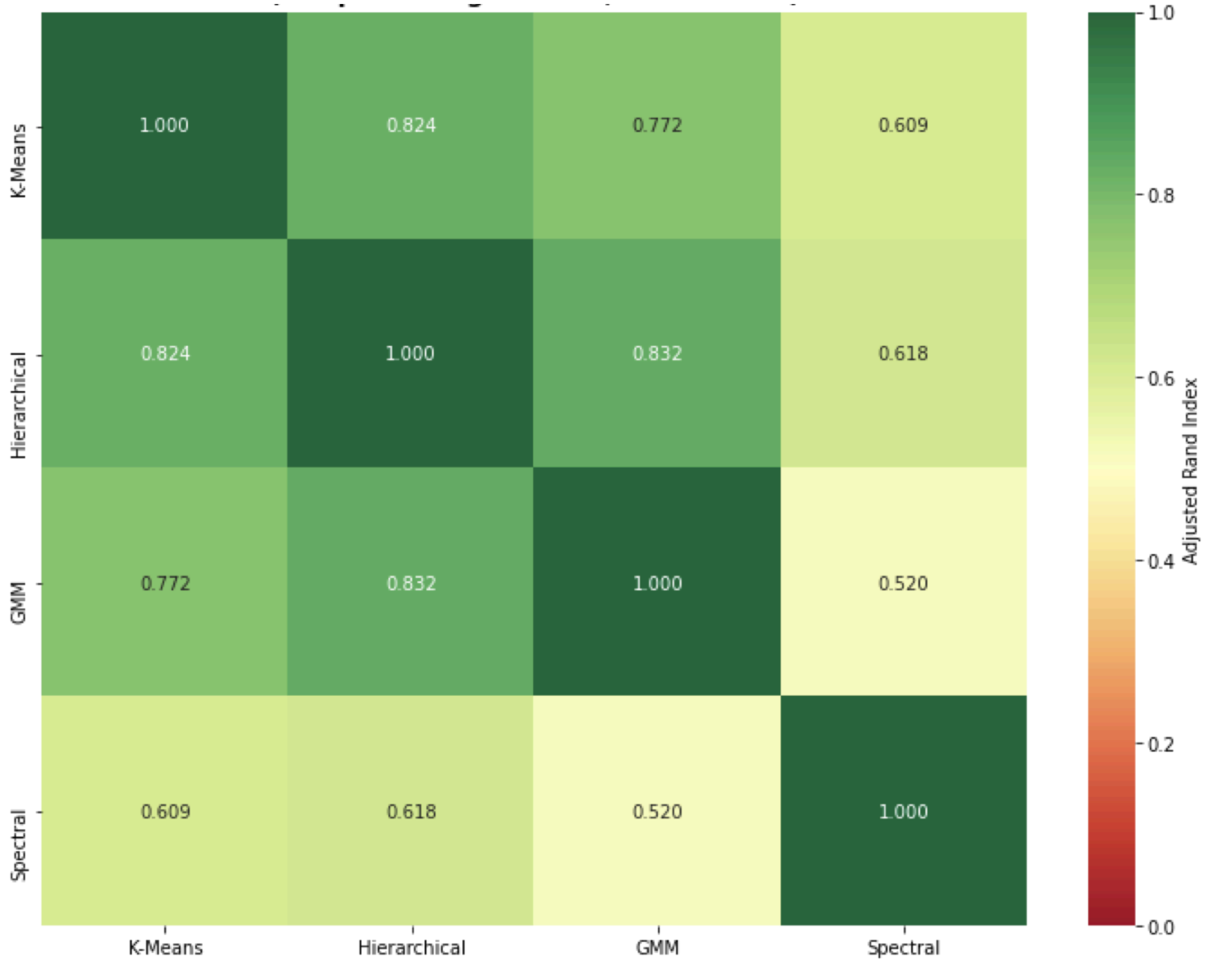


To validate the k-mean clustering results, the 100 control energy trajectories were re-clustered using hierarchical clustering (Ward linkage), Gaussian mixture modeling (GMM), and spectral clustering, each constrained to k = 5, and pairwise agreement between cluster assignments was quantified using the Adjusted Rand Index (ARI; ARI = 1 indicates identical partitions, ARI = 0 indicates chance-level agreement).

**Fig. S5. (Dis)alignment between data-driven clusters and cognitive categories**

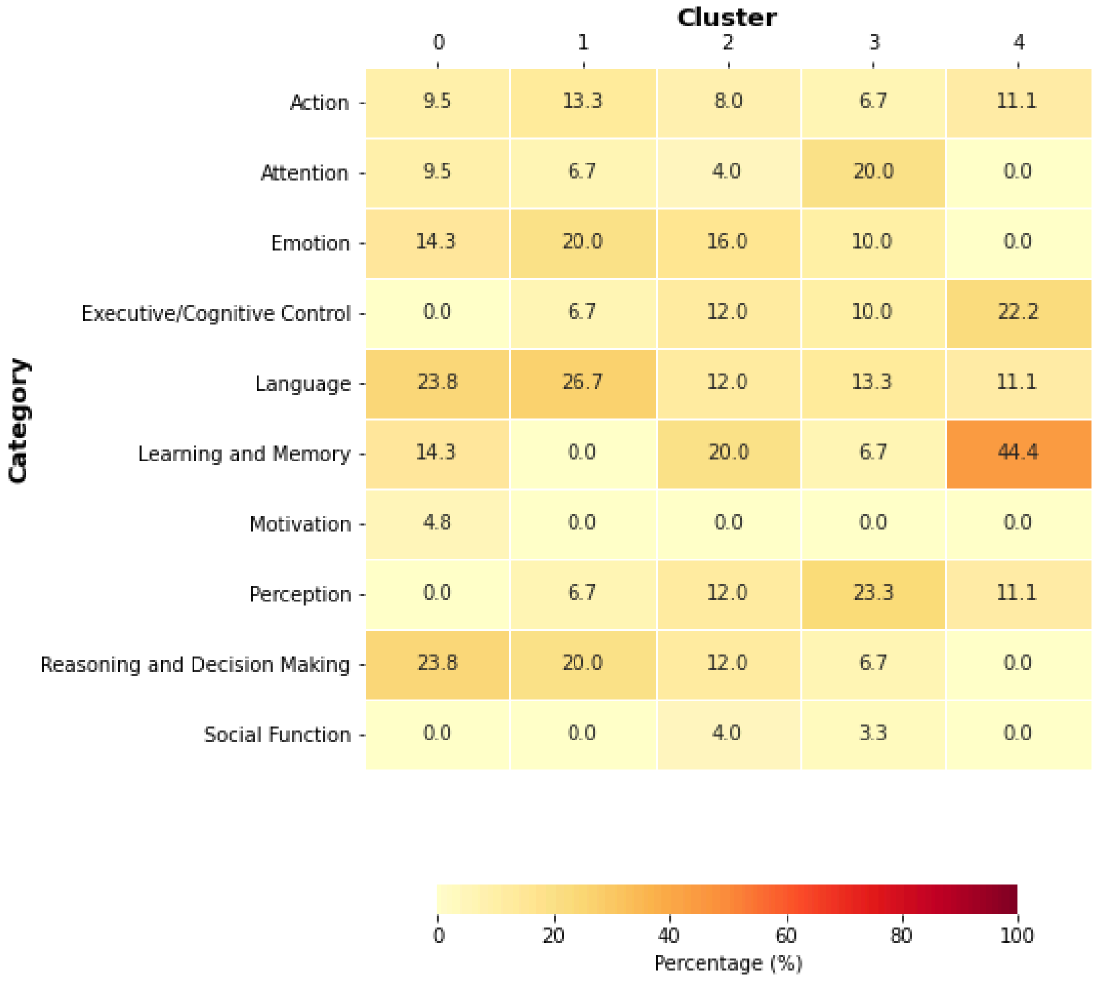


The heatmap shows the percentage of cognitive states (rows) from each Cognitive Atlas Category assigned to each of the data-driven clusters (columns).

**Fig. S6. Transition energy matrix of the control energy cost to switch between all pairs of 100 cognitive states averaged across development.**

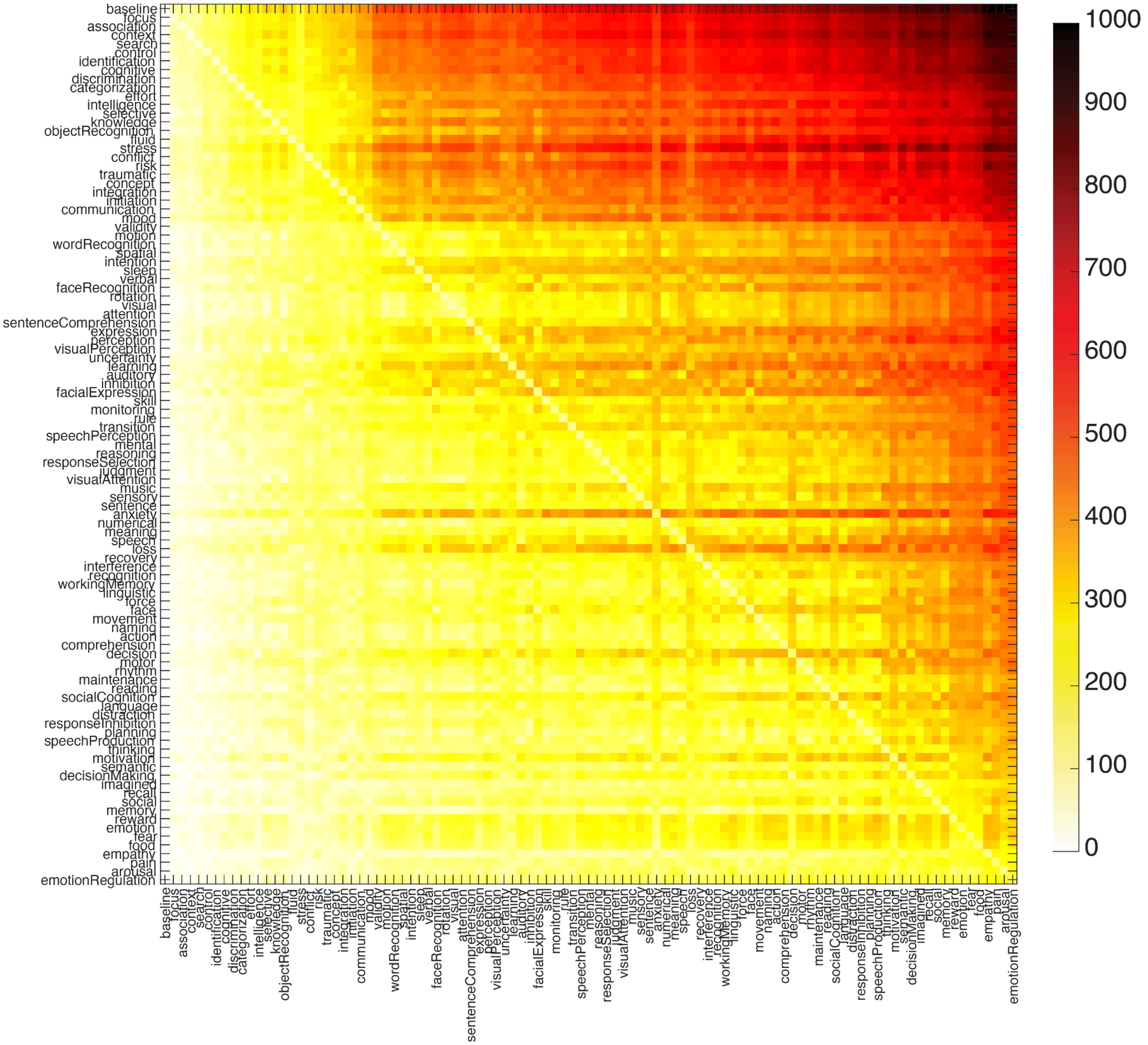

**Table S4. *P*-values of spin-tests against 1,000 spatially autocorrelated null models for each neurodevelopmental event**

| ***p_spin*** | | Myelination | Axon Development | Dendrite Development | Synaptic Development | Neuron Migration | Neuron Differentiation |
|---|---|---|---|---|---|---|---|
| Infant | language | 0.020 | | | | | |
| | reward | | | | | 0.018 | |
| | cognitive | | 0.004 | | | | |
| Toddler | language | | | | | 0.041 | |
| | reading | | | 0.010 | | | |
| | recognition | | | | | 0.028 | 0.029 |
| | verbal | | | | 0.040 | | 0.001 |
| | communication | 0.005 | | | | | |
| Pre-school | language | | | | | | 0.036 |
| | learning | | 0.032 | | | | |
| | recognition | | | | | | |
| | verbal | | | 0.020 | 0.008 | | |
| | auditory | | | | | 0.040 | |
| | social | | | | | | |
| | sleep | | | | | | |
| | motor | 0.006 | | | | | |
| | attention | 0.039 | | 0.042 | | | |
| | memory | | | | | 0.007 | |
| | action | 0.026 | | | | | |
| | cognitive | 0.011 | | | | | |
| | visual | | 0.040 | | | | |
| | communication | | | 0.019 | 0.025 | | |
| School | verbal | | | 0.048 | 0.012 | | |
| | sleep | | 0.044 | | | | |
| | memory | 0.029 | 0.029 | | | | |
| Adolescence | verbal | | | | 0.011 | | |
| | anxiety | 0.038 | | | | | |
| | attention | 0.047 | | 0.031 | | | |
| | memory | | 0.016 | | | | |

**Table S5. T-statistics comparing control energy required to reach vs to leave each cognitive state**

| harder-to-reach states | | | harder-to-leave states | | |
|---|---|---|---|---|---|
| keywords | *t* | *p* | keywords | *t* | *p* |
| discrimination | 2.09 | 0.039 | baseline | -25.47 | 1.66E-45 |
| effort | 5.14 | 1.38E-06 | context | -18.00 | 3.90E-33 |
| knowledge | 11.03 | 5.61E-19 | categorization | -10.04 | 8.25E-17 |
| object recognition | 5.71 | 1.17E-07 | intelligence | -3.66 | 0.00041 |
| fluid | 13.00 | 3.25E-23 | selective | -2.54 | 0.013 |
| conflict | 2.49 | 0.014 | stress | -11.63 | 2.71E-20 |
| traumatic | 8.76 | 5.06E-14 | integration | -3.49 | 0.00072 |
| initiation | 6.32 | 7.42E-09 | communication | -20.64 | 7.97E-38 |
| spatial | 3.43 | 0.00088 | mood | -14.13 | 1.45E-25 |
| intention | 8.74 | 5.58E-14 | validity | -2.96 | 0.0038 |
| sleep | 10.96 | 7.69E-19 | motion | -2.59 | 0.011 |
| face recognition | 4.99 | 2.51E-06 | word recognition | -13.28 | 8.25E-24 |
| rotation | 6.54 | 2.65E-09 | verbal | -14.86 | 4.70E-27 |
| sentence comprehension | 17.00 | 2.87E-31 | visual | -11.20 | 2.42E-19 |
| expression | 3.41 | 0.00094 | attention | -2.84 | 0.0054 |
| perception | 2.30 | 0.024 | learning | -17.92 | 5.53E-33 |
| visual perception | 5.08 | 1.75E-06 | facial expression | -2.20 | 0.030 |
| uncertainty | 10.63 | 4.17E-18 | mental | -3.57 | 0.00054 |
| auditory | 6.85 | 5.97E-10 | response selection | -21.45 | 3.37E-39 |
| inhibition | 3.27 | 0.0015 | sensory | -10.54 | 6.45E-18 |
| monitoring | 3.83 | 0.00023 | anxiety | -15.76 | 7.48E-29 |
| rule | 10.77 | 2.03E-18 | meaning | -18.08 | 2.71E-33 |
| speech perception | 16.92 | 4.16E-31 | speech | -8.28 | 5.66E-13 |
| reasoning | 11.80 | 1.21E-20 | recognition | -13.20 | 1.24E-23 |
| judgment | 7.50 | 2.72E-11 | linguistic | -13.27 | 8.71E-24 |
| visual attention | 17.01 | 2.78E-31 | face | -21.52 | 2.61E-39 |
| music | 5.22 | 9.84E-07 | action | -19.75 | 2.72E-36 |

| | | | | | |
|---|---|---|---|---|---|
| sentence | 13.44 | 3.95E-24 | decision | -11.79 | 1.26E-20 |
| loss | 5.85 | 6.31E-08 | motor | -9.88 | 1.77E-16 |
| recovery | 8.74 | 5.64E-14 | planning | -12.19 | 1.72E-21 |
| interference | 3.56 | 0.00057 | social | -15.91 | 3.72E-29 |
| working memory | 7.63 | 1.41E-11 | emotion | -3.11 | 0.0025 |
| force | 3.70 | 0.00035 | food | -12.95 | 4.20E-23 |
| movement | 11.82 | 1.08E-20 | empathy | -7.85 | 4.72E-12 |
| naming | 11.57 | 3.79E-20 | | | |
| rhythm | 4.69 | 8.68E-06 | | | |
| maintenance | 13.86 | 5.19E-25 | | | |
| language | 8.25 | 6.41E-13 | | | |
| distraction | 6.32 | 7.49E-09 | | | |
| speech production | 17.34 | 6.52E-32 | | | |
| thinking | 8.82 | 3.73E-14 | | | |
| motivation | 4.86 | 4.43E-06 | | | |
| semantic | 6.62 | 1.82E-09 | | | |
| decision making | 2.73 | 0.0075 | | | |
| recall | 4.21 | 5.60E-05 | | | |
| memory | 12.15 | 2.09E-21 | | | |
| fear | 9.52 | 1.14E-15 | | | |
| pain | 14.28 | 7.06E-26 | | | |
| arousal | 10.73 | 2.53E-18 | | | |

**Fig. S7. Developmental stability of the transition matrix across all developmental stages**

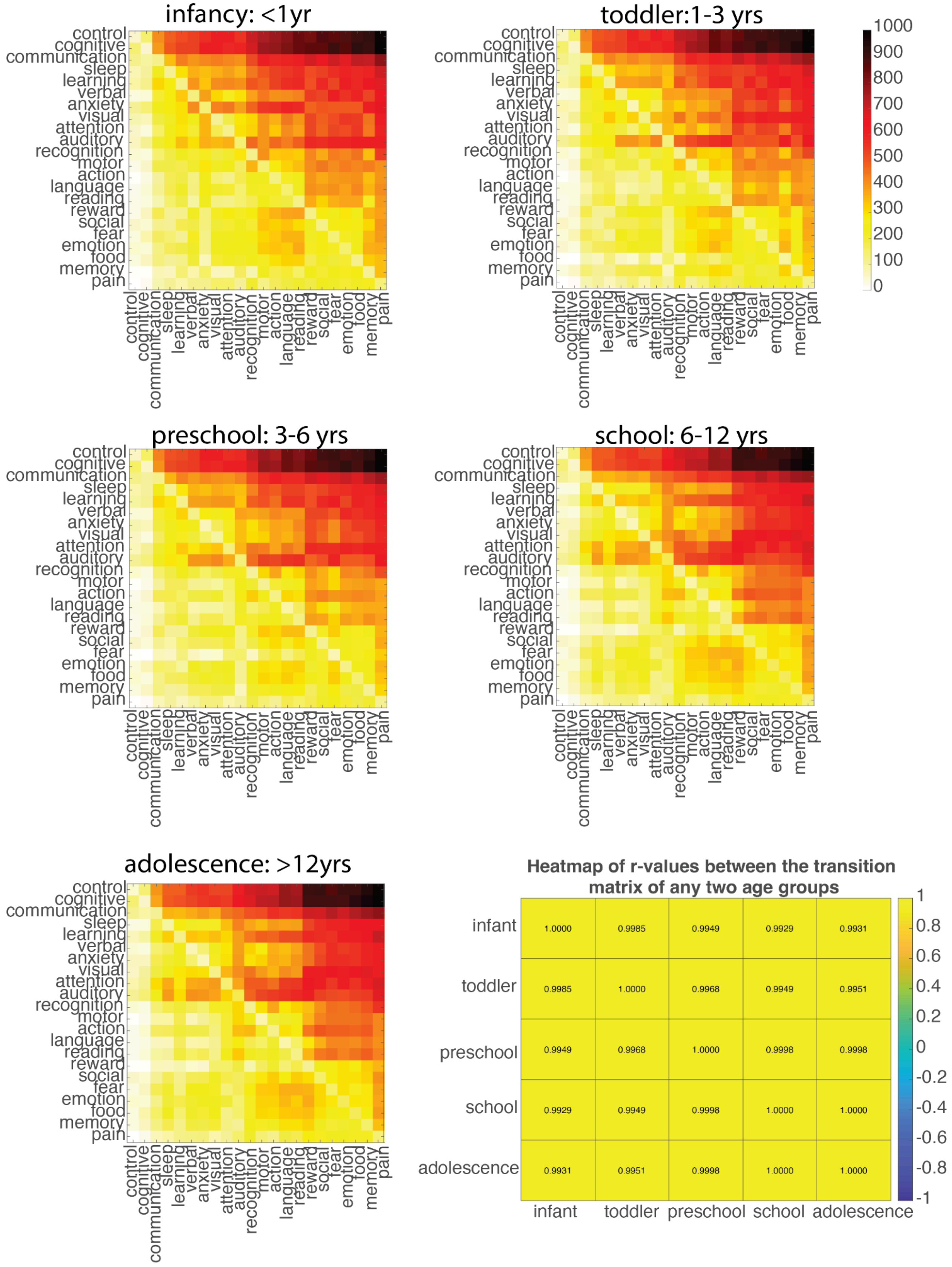

**Fig. S8. T-statistic comparing within-category versus cross-category transition costs over age.**

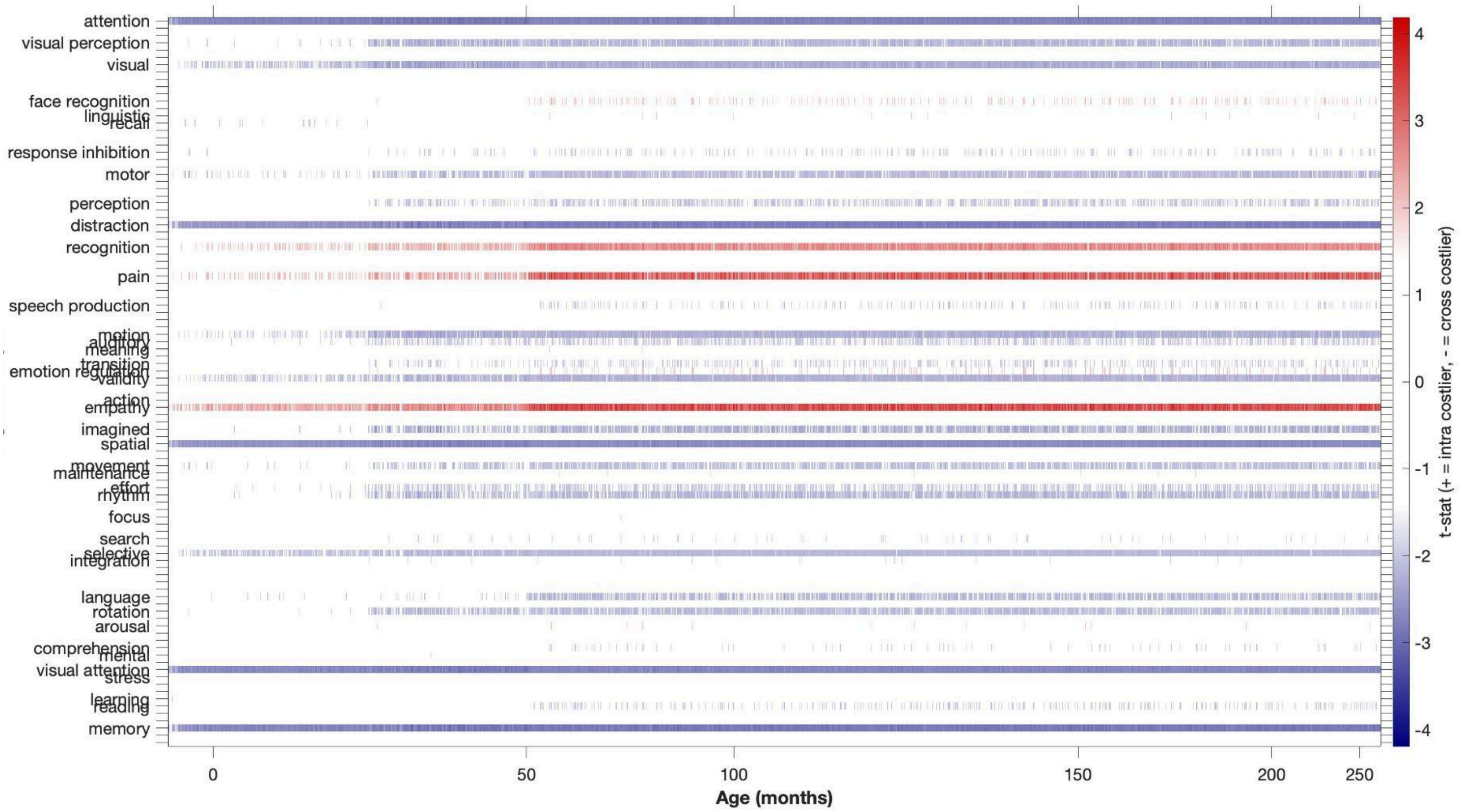


The heatmap displays t-statistics from pairwise comparisons of within-category versus cross-category control energy at each age (x-axis, in months) for each cognitive keyword (y-axis), with only statistically significant values shown. Positive values (red) indicate that within-category transitions are costlier than cross-category transitions; negative values (blue) indicate the opposite pattern.

**Fig. S9. Developmental trajectories and growth rate curves of control energy for the 11 overlapping cognitive states defined by BrainMap**

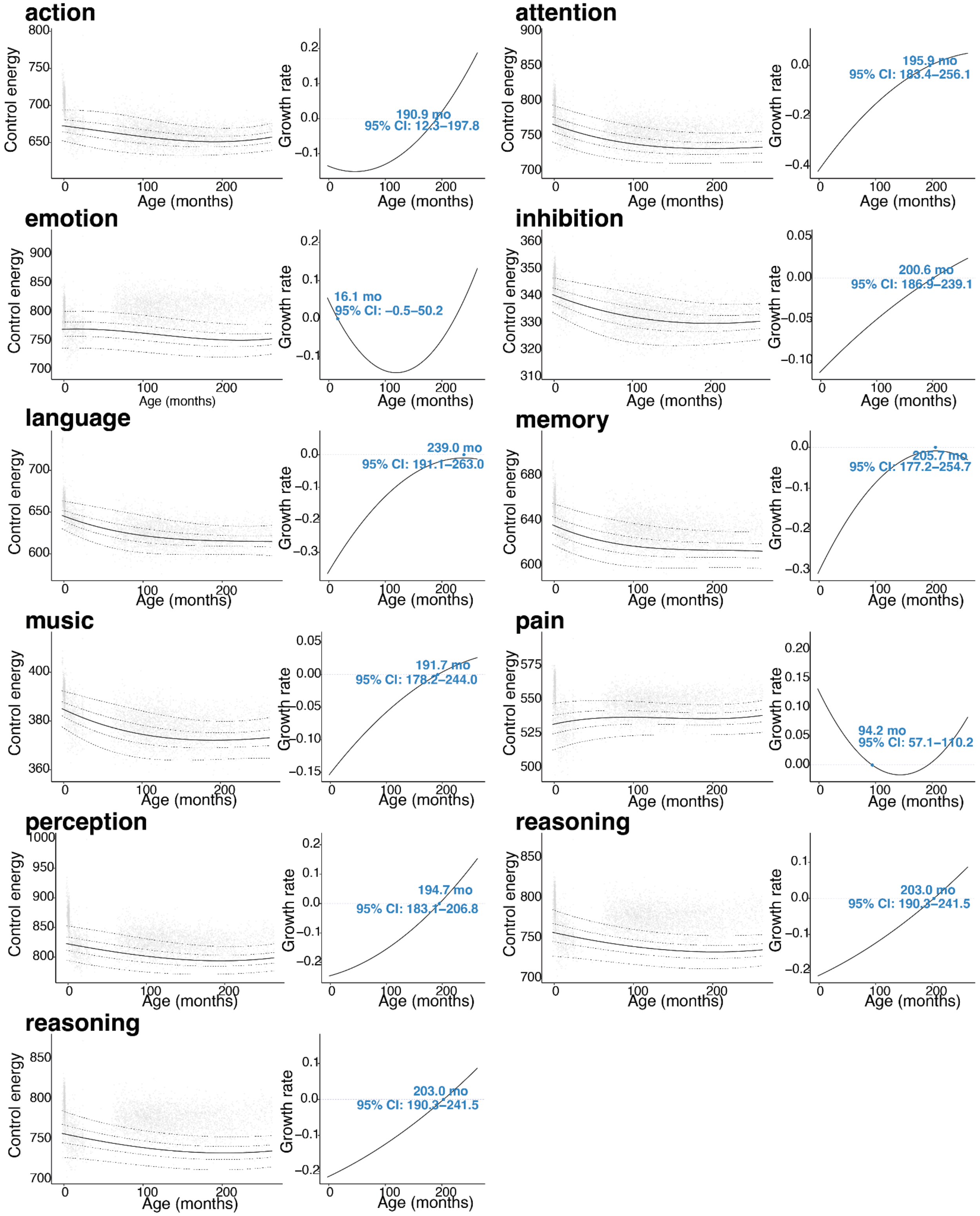

### Supplementary methods

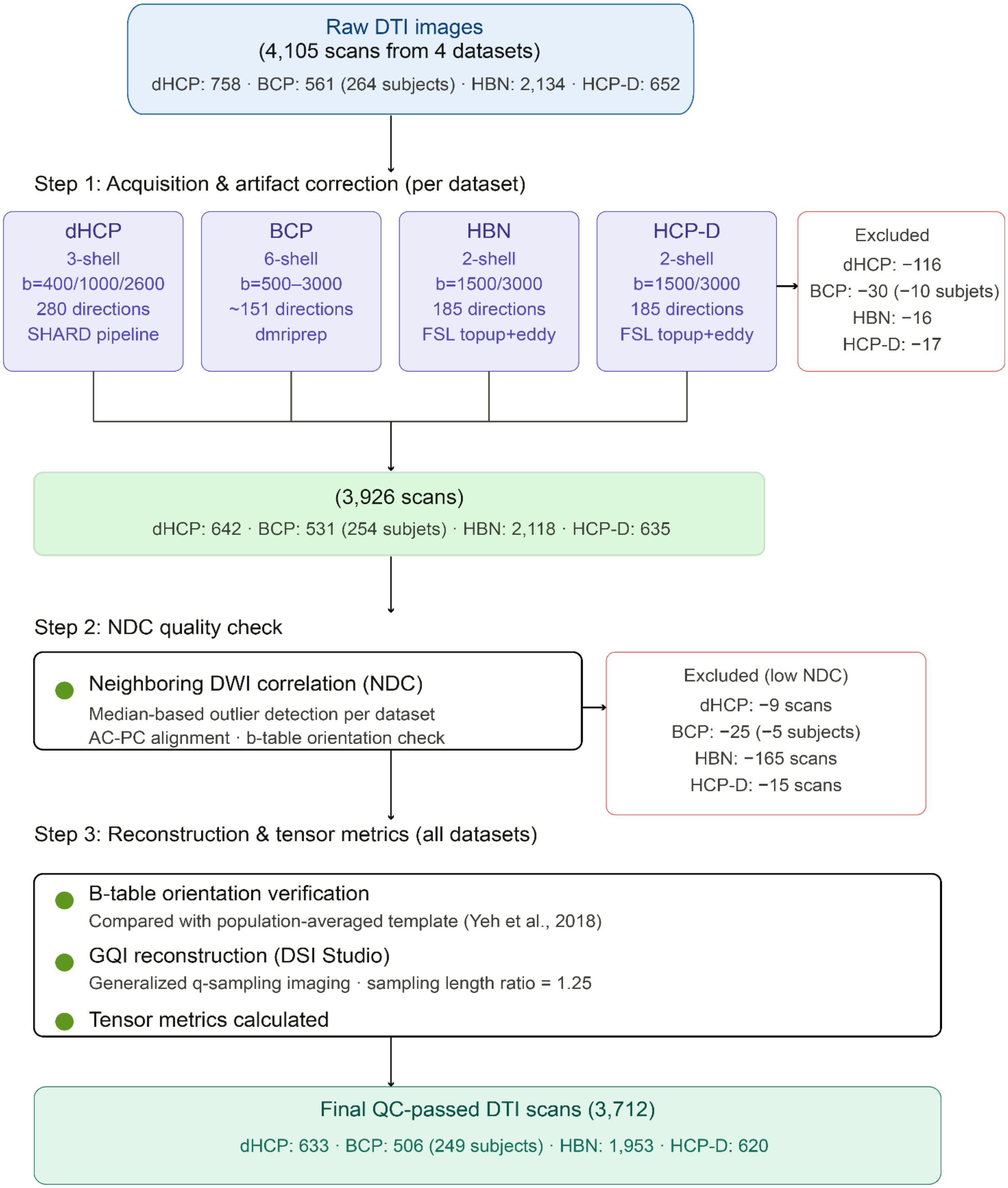


**Fig. S10. DTI data quality-control and preprocessing pipeline across four developmental neuroimaging datasets.** The whole quality-control process resulted in the total exclusion of 393 scans. The final sample included 3,712 scans (dHCP: 633; BCP: 506 from 249 participants; HBN: 1,953; HCP-D: 620), spanning 27 postmenstrual weeks to 22 years of age.

**DTI data preprocessing for each dataset**

**dHCP:** A multishell diffusion scheme was used, and the b-values were 400, 1000, and 2600 s/mm². The number of diffusion sampling directions was 64, 88, and 128, respectively. The in-plane resolution was 1.5 mm. The slice thickness was 1.5 mm. The images were denoised and corrected for Gibbs ringing, motion, eddy current, and susceptibility artifact using the diffusion SHARD pipeline. A quality check was conducted using neighboring DWI correction (NDC) (Yeh, NeuroImage, 2019 Nov 15;202:116131). 125 out of 758 scans (including repeated scans) were excluded: 116 due to failure at one or more steps of the processing pipeline, and a further 9 due to low NDC values identified by a median-based outlier detector. The accuracy of b-table orientation was examined by comparing fiber orientations with those of a population-averaged template (Yeh et al., NeuroImage, 2018). The restricted diffusion was quantified using restricted diffusion imaging (Yeh et al., MRM, 77:603–612, 2017). The diffusion data were reconstructed using generalized q-sampling imaging (Yeh et al., IEEE TMI, 29(9):1626–35, 2010) with a diffusion sampling length ratio of 1.25. The tensor metrics were calculated. The analysis was conducted using the resource allocation (TG-CIS200026) at Extreme Science and Engineering Discovery Environment (XSEDE) resources (Towns, J. et al., Computing in Science & Engineering, 16, 62–74, 2014).

**BCP**: The BCP dataset uses a multi-shell protocol with b-values of 500, 1000, 1500, 2000, 2500, and 3000 s/mm², totaling around 151 diffusion directions. The in-plane resolution was 1.5 mm. The slice thickness was 1.5 mm. Artifacts were addressed by dmriprep, the officially recommended dMRI processing pipeline for BCP, and included eddy currents, head motion, bed vibration and pulsation, venetian blind artifacts, slice-wise and gradient-wise intensity inconsistencies, and susceptibility artifacts. 55 out of 561 scans (from 264 participants) were excluded: 30 due to failure at one or more steps of the processing pipeline (reducing the participant count to 254), and a further 25 due to low NDC values identified by a median-based outlier detector (reducing the participant count to 249), yielding a final sample of 506 scans from 249 participants. The accuracy of b-table orientation was examined by comparing fiber orientations with those of a population-averaged template (Yeh et al., NeuroImage, 2018). The restricted diffusion was quantified using restricted diffusion imaging (Yeh et al., MRM, 77:603–612, 2017). The diffusion data were reconstructed using generalized q-sampling imaging (Yeh et al., IEEE TMI, 29(9):1626–35, 2010) with a diffusion sampling length ratio of 1.25. The tensor metrics were calculated. The analysis was conducted using the resource allocation (TG-CIS200026) at Extreme Science and Engineering Discovery Environment (XSEDE) resources (Towns, J. et al., Computing in Science & Engineering, 16, 62–74, 2014).

**HBN/HCPD**: A multishell diffusion scheme was used, and the b-values were 1500 and 3000 s/mm². The number of diffusion sampling directions was 93 and 92, respectively. The in-plane resolution was 1.5 mm. The slice thickness was 1.5 mm. The susceptibility and eddy current artifacts were corrected using FSL topup and eddy (FMRIB, Oxford), implemented through the integrated interface in DSI Studio ("Chen" release). The diffusion MRI data were rotated to align with the AC-PC line. For HBN, 181 out of 2,134 scans were excluded: 16 due to failure at one or more steps of the processing pipeline, and a further 165 due to low NDC values, yielding a final sample of 1,953 scans. For HCP-D, 32 out of 652 scans were excluded: 17 due to failure at one or more steps of the processing pipeline, and a further 15 due to low NDC values, yielding a final sample of 620 scans. The accuracy of b-table orientation was examined by comparing fiber orientations with those of a population-averaged template (Yeh et al., NeuroImage, 2018). The restricted diffusion was quantified using restricted diffusion imaging (Yeh et al., MRM, 77:603–612, 2017). The diffusion data were reconstructed using generalized q-sampling imaging (Yeh et al., IEEE TMI, 29(9):1626–35, 2010) with a diffusion sampling length ratio of 1.25. The tensor metrics were calculated. The analysis was conducted using the resource allocation (TG-CIS200026) at Extreme Science and Engineering Discovery Environment (XSEDE) resources (Towns, J. et al., Computing in Science & Engineering, 16, 62–74, 2014).

**GAMLSS modeling**

<u>Family distribution selection</u>

To select an appropriate distributional family, we fitted GAMLSS models with 10 candidate families — Normal (NO), Student's t (TF), Power Exponential (PE), Log-Normal (LOGNO), Gamma (GA), Inverse Gaussian (IG), Box-Cox t (BCT), Box-Cox Power Exponential (BCPE), Johnson's SU (JSU), and Skew t type 3 (ST3) — to each of the 100 node energy outcomes. BCT achieved the lowest AIC in 42 outcomes and was within two AIC units of the best-fitting family in 62 outcomes, representing the most consistently well-fitting family across outcomes. BCT was therefore adopted as a uniform distributional family to facilitate parameter comparability across nodes.

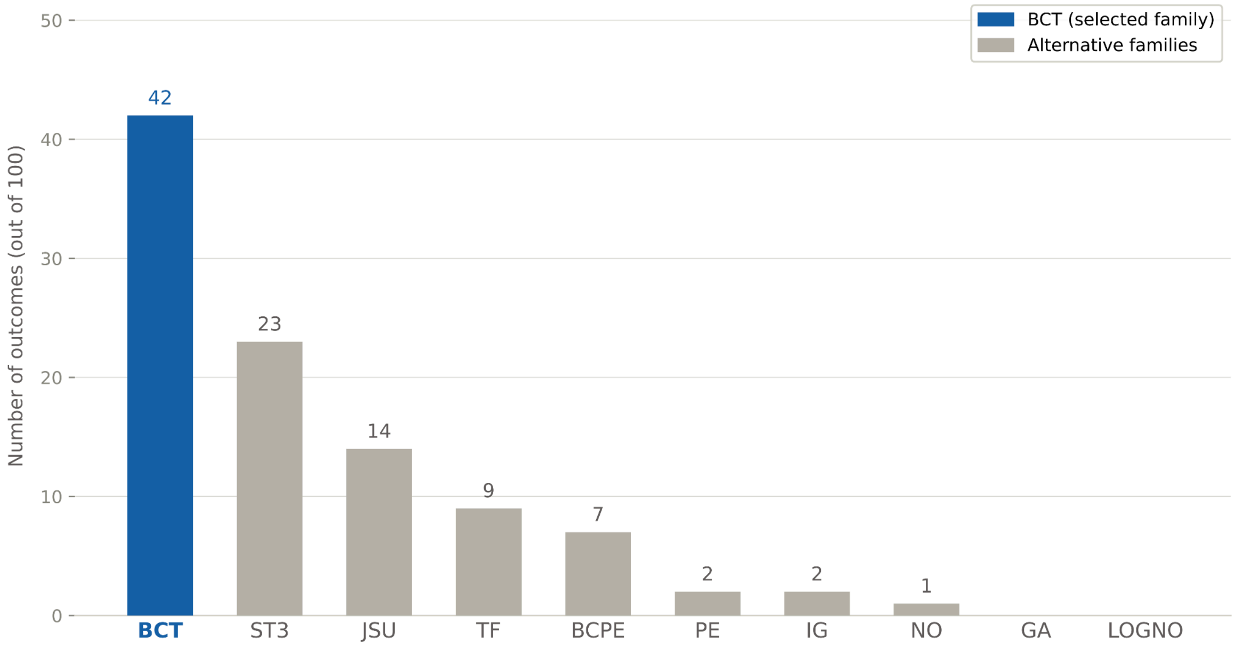


**Fig. S11. Distribution family selection for GAMLSS models across the control energy trajectories of the 100 cognitive states.** Bars indicate the number of outcomes for which each of 10 candidate distribution families achieved the lowest Akaike Information Criterion (AIC). The Box-Cox t (BCT) family achieved the best fit in 42 of 100 outcomes and was within $\Delta AIC \leq 2$ of the best-fitting family in 62 outcomes, representing the most consistently well-fitting family overall. BCT was therefore adopted as a uniform distributional family to fit all control energy changes for all 100 cognitive states to facilitate comparability of model parameters. (BCT = Box-Cox t; ST3 = Skew t Type 3; JSU = Johnson's SU; TF = Student's t; BCPE = Box-Cox Power Exponential; PE = Power Exponential; IG = Inverse Gaussian; NO = Normal; GA = Gamma; LOGNO = Log-Normal.)
